# Conserved assembly architecture of the essential herpesvirus packaging accessory factor

**DOI:** 10.64898/2026.01.22.701024

**Authors:** Elizabeth J. Bailey, Swapnil C. Devarkar, Renata Szczepaniak, Laura M. Meißner, Xinyu Chen, Chunxiang Wu, Sandra K. Weller, Yong Xiong, Allison L. Didychuk

## Abstract

To create a new wave of infectious virions, all herpesviruses require an accessory factor of unknown function to package their viral genomes into nascent capsids. Here, we present cryo-EM structures of the packaging accessory factor from the α-herpesvirus herpes simplex virus type 1 (HSV-1, UL32) and the β-herpesvirus human cytomegalovirus (HCMV, UL52). Unlike homologs from the γ-herpesviruses, neither UL32 nor UL52 form stable homopentameric rings. UL52 forms incomplete pentameric rings lacking one or two protomers. UL32 does not form stable higher-order species, but stabilization through chemical crosslinking revealed a novel quaternary structure where three pentameric rings assemble into a “tripentamer.” Our results reveal that herpesvirus packaging accessory factors adopt distinct oligomeric states but are constrained to pentameric symmetry. Assembly of protomers into a ring creates a positively charged central channel that we show is critical for infectious virus production in HSV-1. Taken together, our study points to a structurally conserved, essential function of packaging accessory factors across the *Herpesviridae*.

**AUTHOR SUMMARY:** Herpesviruses are a diverse family of pathogens that drive human disease. At the end of their life cycle, all herpesviruses must actively package their genomes into newly formed capsids. Although some aspects of packaging are conserved with tailed bacteriophages, herpesviruses require an additional packaging accessory factor of unknown function. Here, we present two high-resolution structures of the packaging accessory factor from herpes simplex virus and human cytomegalovirus and identify pentameric symmetry as a unifying structural feature. We propose that the packaging accessory factor oligomerizes to concentrate positively charges in a central channel critical for packaging progression.

## INTRODUCTION

Nearly every human is infected with at least one herpesvirus by adulthood. Nine herpesviruses from three subfamilies (α, β, γ) infect humans and cause clinically significant disease. Herpes simplex virus type 1 (HSV-1) is an α-herpesvirus that causes oral and genital lesions and can lead to disseminated disease in neonates^1^. The β-herpesvirus cytomegalovirus (HCMV) is the leading non-genetic cause of congenital birth defects^2^ and causes complications in transplant patients. Kaposi’s Sarcoma-associated herpesvirus (KSHV) is a γ-herpesvirus that causes cancer and other malignancies, particularly in immunocompromised individuals^3,4^.

Late in their replication cycle, herpesviruses compress their double-stranded DNA genome into capsids that mature to form infectious progeny virions^5^. Empty icosahedral capsids, with a unique portal vertex through which DNA is inserted, are assembled in the nucleus of infected cells^6–10^. Genome packaging is driven by the virus-encoded terminase motor, which recognizes newly synthesized viral genomes, translocates the viral genome into the nascent capsid, and finally cleaves the packaged, unit-length viral genome^11–13^. The genome-filled capsid is stabilized by binding of the portal cap, followed by egress of the capsid through the nuclear membrane to continue maturation^14–17^.

While this packaging process is largely conserved with tailed bacteriophages^18,19^, herpesviruses require an additional packaging accessory factor of unknown function. This factor is unique to herpesviruses and essential for viral genome packaging across the *Herpesviridae*^20–25^. The viral genome is replicated in the absence of the packaging accessory factor, but the concatenated genome is not cleaved, consistent with a defect early in packaging^20,22,23^. Loss of the packaging accessory factor results in immature “B” capsids that retain the cleaved scaffold and lack packaged DNA^20,22–24^. The packaging accessory factor is not incorporated into capsids or virions^20,26^, nor is it thought to be a constitutive component of the terminase, although a recent study in HCMV suggests that the packaging accessory factor and terminase may interact^27^.

The KSHV packaging accessory factor, ORF68, adopts a novel fold and oligomerizes into a stable homopentameric ring with a positively charged central channel that is essential for production of infectious virions^28^. While homologs from related γ-herpesviruses can partially complement for loss of ORF68 during KSHV infection, packaging accessory factors from the more distantly related α- and β-herpesviruses cannot^28^. Thus, we sought to determine if the packaging accessory factor is structurally conserved across the herpesvirus family. Here, we report cryo-electron microscopy (cryo-EM) structures of the packaging accessory factor from HSV-1 (UL32) and HCMV (UL52). We find that individual protomers adopt a highly conserved core tertiary structure, but their propensity to form stable oligomers varies across the *Herpesviridae*. HCMV UL52 assembles into incomplete pentameric rings with three or four protomers and a slight helical twist. HSV-1 UL32 formed an unstable pentamer in solution, leading us to pursue chemical crosslinking that stabilized a novel quaternary assembly composed of three pentameric rings. Assembly into a ring concentrates positively charged residues into a central channel, and we demonstrate that these positively charged residues are essential to productive infection in HSV-1. Our structures of UL52 and UL32 form the basis for further mechanistic dissection of this essential yet enigmatic viral protein.

## RESULTS

### The conserved herpesvirus packaging accessory factor adopts different oligomeric states across the *Herpesviridae*

HSV-1 UL32 and HCMV UL52 have insertions and amino (N)-terminal extensions relative to KSHV ORF68, resulting in protomers that are 12 and 22 kDa larger than ORF68 (**Figs 1a**, **S1a**). Recombinantly expressed ORF68, UL32, and UL52 were purified to homogeneity from baculovirus-infected insect cells (**Fig 1b**). During size exclusion purification, we noticed that ORF68 eluted as a single peak at a volume consistent with a pentameric assembly, while UL32 and UL52 eluted later than expected for pentameric assemblies. To monitor oligomeric states in solution, we performed size exclusion chromatography coupled with multi-angle light scattering (SEC-MALS) on purified samples of the three packaging accessory factors. ORF68 eluted in a single peak with a calculated molecular weight of ∼280 kDa, consistent with a pentamer (**Fig 1c**, **1d**)^28^. In contrast, UL52 exhibited a broad peak with a molecular weight corresponding to between 1-2 protomers and UL32 had two well-separated peaks consistent with the molecular weights of a pentamer and a monomer (**Fig 1c**, **1d**). Thus, unlike the γ-herpesvirus accessory factors ORF68 and the Epstein-Barr virus (EBV) homolog BFLF1^28^, UL32 and UL52 do not form stable pentameric rings in solution. We next sought to structurally characterize UL32 and UL52 to illuminate the tertiary structure and quaternary assemblies of these essential proteins.

**Fig 1.**
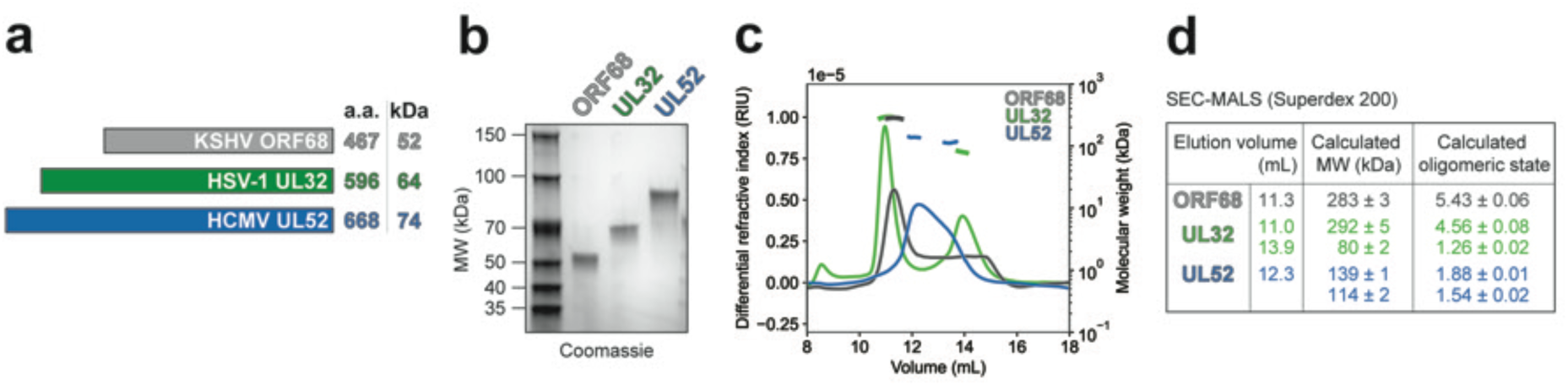
ORF68 homologs adopt different quaternary states in solution. (a) Packaging accessory factors from KSHV (ORF68), HSV-1 (UL32), and HCMV (UL52) range in size from 52 kDa to 74 kDa. (b) SDS-PAGE gel of purified ORF68, UL32, UL52. (c) SEC-MALS traces for ORF68, UL32, UL52. (d) Table of SEC-MALS elution volumes and calculated oligomeric states.

### UL52, the HCMV packaging accessory factor, forms an incomplete pentameric ring

We carried out structural studies of UL52 using single-particle cryo-EM and identified two classes of incomplete pentameric rings with three or four well-resolved protomers (**Figs 2a**, **S2, S1 Movie, Table 1**). We were unable to identify any particles with complete pentameric rings. The maps for the 3-membered incomplete ring and the 4-membered incomplete ring each had a global resolution of 3.29 Å (hereafter referred to as 3-mer and 4-mer, respectively). The UL52 model was first built for the protomer with the highest local resolution and was used as a starting model to build the other protomers and create the final 4-mer model (**Figs 2b**, **S2e**). In our processing we also saw a class of 2-mer particles (**Fig S2a**) suggesting that UL52 can form oligomers of 2, 3, and 4 protomers.

**Fig 2.**
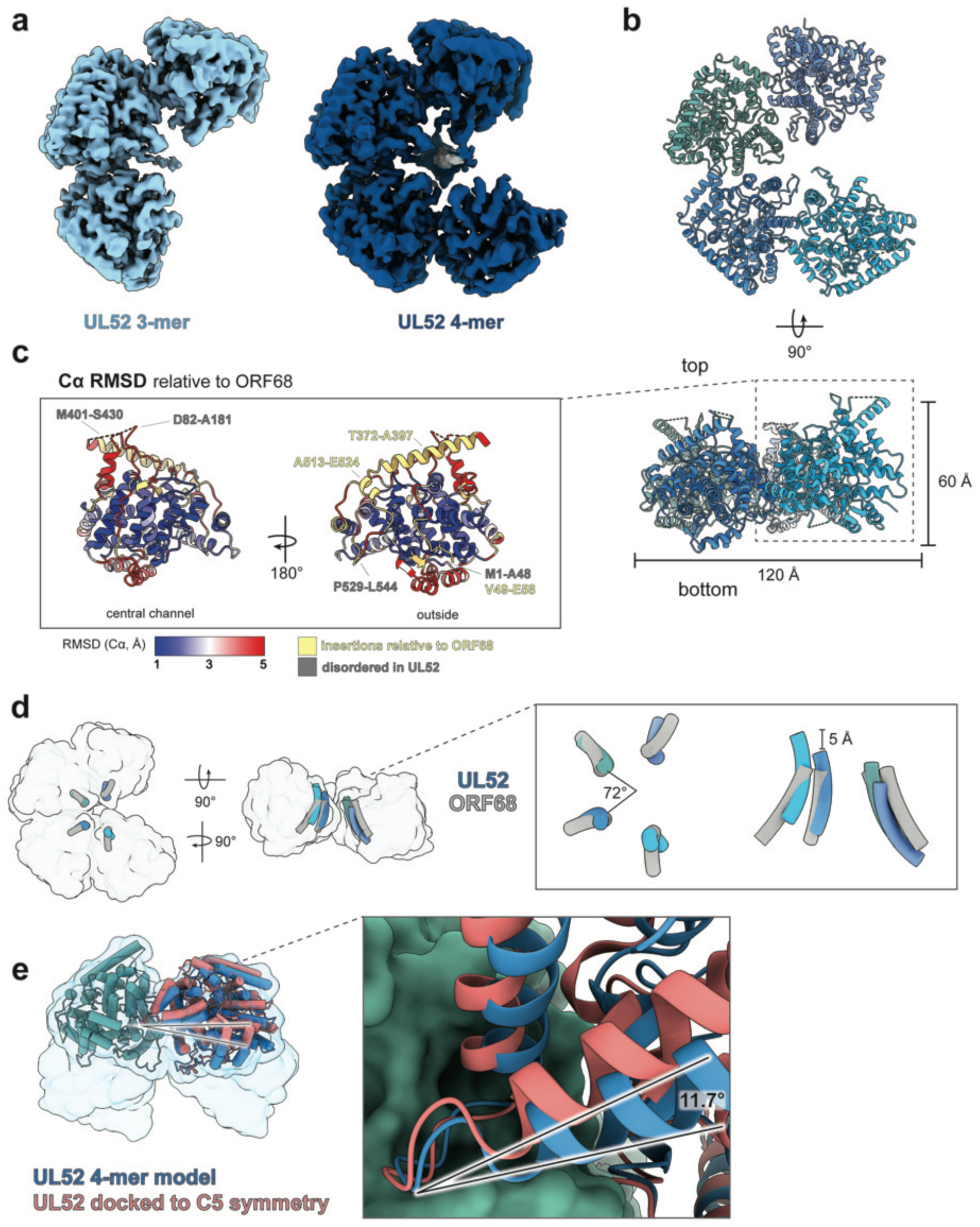
HCMV UL52 forms an incomplete ring. (**a**) Cryo-EM densities of the UL52 3-membered incomplete ring (3-mer, light blue, 119 k particles, C1, 3.29 Å) and 4-membered incomplete ring (4-mer, dark blue, 147 k particles, C1, 3.29 Å) colored by proximity to the models. Unmodeled density (grey) is resolved in the central channel. (**b**) Model of the UL52 4-mer shown from top and side views, with each protomer colored a shade of blue. (**c**) UL52 protomer colored by the Cα RMSD to ORF68 (PDB ID: 6XF9), blue is 1 Å, white is 3 Å, and red is 5 Å, yellow are residues without RMSD values where there are insertions in UL52 relative to ORF68. (**d**) UL52 adopts pentamer-like symmetry with a twist out of plane. Central channel helices for ORF68 (grey) and UL52 are aligned to a UL52 protomer (sage green). Models are shown in tube helices inside transparent UL52 4-mer model surface at 10 Å resolution. Insert: (Left) shows a top view of the 5-fold axis (72°, helical twist (Δ*ϕ)*). (Right) shows a side view of the right-handed twist of the UL52 protomers relative to the planar arrangement of ORF68 and the ∼5 Å helical rise (Δ*z*) between protomers. (**e**) UL52 twists at the protomer interface loop. UL52 protomer models (sage, blue) shown relative to a UL52 protomer docked to a hypothetical C5 planar arrangement (salmon). Models are shown within a transparent UL52 4-mer model surface at 10 Å resolution. Inset shows details of the protomer interface pocket/loop region, where the pocket protomer model is shown as a sage surface and the loop protomer models are shown as blue or salmon cartoons.

**Table 1.**
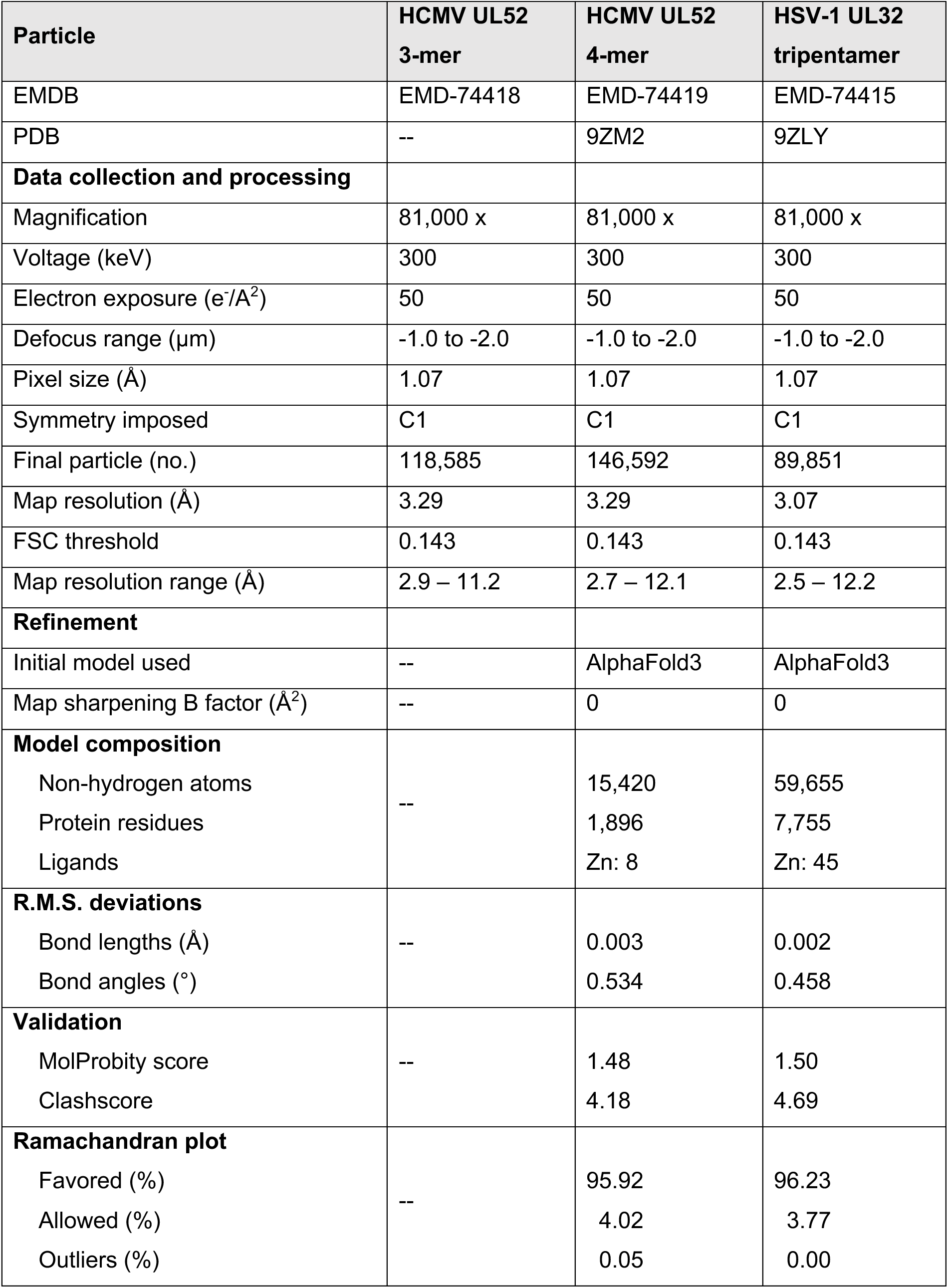
Cryo-EM data collection, refinement, and validation statistics.

For the purposes of our discussion, we define the orientation of packaging accessory factors based on the crown-like ring in Didychuk *et al.*^28^, where the “top” surface of the ring is crenellated by short semi-structured loops and the “bottom” surface of the ring is relatively flat. The core fold of UL52 is highly similar to that of ORF68 (**Fig 2c**). The outer surface is more variable, with long insertions that form surface-exposed helices and loops and an N-terminal extension. The 51-amino acid N-terminal extension is almost entirely disordered, with residues 1-48 unresolved (**Figs 2c**, **S3a, S3b**). Also unresolved are two long, negatively charged insertions on the top surface of the incomplete ring (residues 82-181, 401-430) (**Figs 2c**, **S3a, S3b**). Both the 3-mer and 4-mer cryo-EM classes have additional unidentified low-resolution density in the central channel (**Fig 2a**), which could be attributed to co-purified nucleic acid or part of the unresolved, negatively charged loop (residues 82-181) interacting with the positively charged central channel.

UL52 has two regions that differ relative to ORF68, altering the size and shape of the top and side of the UL52 incomplete ring. A long α-helix insertion and subsequent 30 amino acid disordered loop (region 1: residues 372-432) reshape the top peripheral surface of the incomplete ring (**Figs 2c**, **S3a, S3b**). A short α-helical insertion and a 16 amino acid disordered loop (region 2: residues 502-547) reshape the side, peripheral surface (**Figs 2c**, **S3a, S3b**). A sequence alignment of packaging accessory factors from the nine human herpesviruses revealed that the insertion forming UL52 region 2 is unique to β-herpesvirus factors (**Fig S1a**). These differences in the exterior of the protein alter the available surface for binding partners, potentially enabling virus-specific interactions.

The UL52 3-mer and 4-mer both form incomplete rings with pentamer-like symmetry wherein the protomers are arranged ∼72° from each other. However, the incomplete ring twists out of plane, creating a right-handed screw axis with a helical rise of ∼5 Å per protomer (**Fig 2d**). This slight rise is insufficiently steep for UL52 to form a helical filament. In this helical arrangement UL52 protomers adopt a different relative conformation than they would as a planar ring. Examination of the protomer interface reveals that the twist arises from a ∼12° rotation from the helix-loop-helix that extends into a pocket on the adjacent protomer (residues 487-497, hereafter referred to as the protomer interface loop) (**Fig 2e**). The protomer interface loop has several residues buried deep in the pocket of the adjacent protomer, including N489 and K491, as well as polar-charged interactions between loop-side R495 and pocket-side Q286 (**Fig S2g**). Several residues in the loop-pocket regions of the protomer interface are universally- or well-conserved across herpesviruses (**Figs 2f**, **S1b, S1c**). A previous study showed that an insertion in the protomer interface loop of the HSV-1 homolog UL32 was not tolerated^29^. Thus, this loop is the conserved “key” that fits into the “lock” of the adjacent protomer, controlling oligomerization of the packaging accessory factor.

### UL32, the HSV-1 packaging accessory factor, forms an unstable pentameric ring that can be stabilized in a novel quaternary assembly

Like HCMV UL52, SEC-MALS of the HSV-1 packaging accessory factor UL32 revealed multiple oligomeric states. In contrast to the poorly defined UL52 oligomer, we observed two well-separated peaks for UL32 consistent with monomer and pentamer species (**Fig 1c**, **1d**). We first attempted to visualize the pentameric species by negative stain electron microscopy but were unable to identify sufficient particles of the expected dimensions. We suspect that UL32 forms an unstable pentamer at higher concentrations that is disrupted at lower concentrations. We previously reported that ORF68 binds double-stranded DNA (dsDNA) *in vitro*^28^ and hypothesized that if UL32 similarly bound dsDNA, we could use this approach to stabilize an oligomeric complex. We used an electrophoretic mobility shift assay (EMSA) to monitor binding of UL32 to fluorescently labeled dsDNA. UL32 bound to all probes tested, including the shortest (10 bp, **Fig S4a**). The migration of this complex was comparable to ORF68-probe complexes^28^, suggestive of UL32 binding as a pentamer. As the probe length increased to 30 bp, a larger discrete complex formed (**Fig S4a**). UL32-DNA complexes were unstable on cryo-EM grids and thus we pursued chemical crosslinking with ethylene glycol bis(succinimidyl succinate) (EGS), which yielded distinct higher molecular weight bands by SDS-PAGE (**Figs S4b, S4c**). By negative stain electron microscopy, crosslinked samples of UL32 lacking dsDNA or with probes of 10, 20, 30, or 67 bp showed well-dispersed particles, including monomers, pentamers, and a larger species (**Fig S4d**). We anticipated that the larger species would be a dimer of homopentamers (D5 symmetry) as previously observed for the EBV packaging accessory factor BFLF1^28^.

We imaged samples of UL32 crosslinked in the presence of 30 bp dsDNA by single-particle cryo-EM and generated a map with a global resolution of 3.09 Å (**Figs 3a**, **S5, S2 Movie, Table 1**). Our cryo-EM reconstruction of UL32 revealed a novel quaternary assembly wherein three pentameric UL32 rings form a trimer of homopentamers, hereafter referred to as a “tripentamer”. No symmetry was applied during reconstruction (C1). Within the tripentamer, UL32 forms a pentameric planar ring with similar dimensions to those of the ORF68 pentamer, where the top of each pentameric ring faces outward (**Fig 3b**). Like UL52, the core fold of UL32 is similar to that of ORF68 (**Fig 3c**). Compared to ORF68, the surface of UL32 is altered by structured insertions (residues 315-322, 353-390), disordered loops (residues 64-106, 227-234, 265-274), and a partially disordered N-terminal extension (residues 1-31) (**Figs 3c**, **S5g**). A long α-helical insertion (residues 353-390) and two disordered loops (residues 64-106, 265-274) are present in UL32 at similar locations to these features in UL52. Residues 315-322 extend an α-helix on the bottom surface of the ring (residues 308-323) that is displaced laterally by 12 Å relative to the corresponding helices in UL52 and ORF68 (**Fig S5h**).

**Fig 3.**
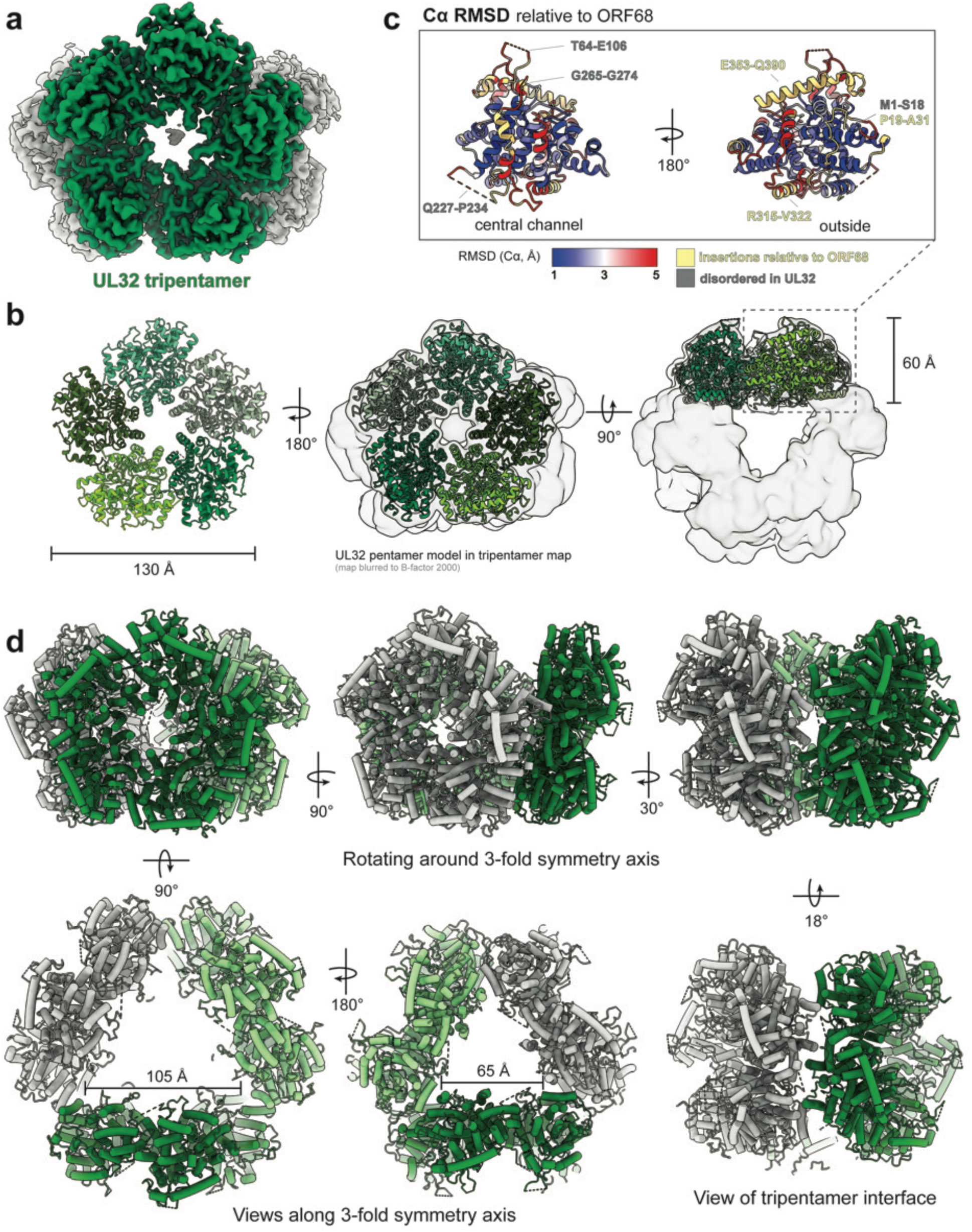
HSV-1 UL32 adopts a novel tripentameric architecture. (**a**) Cryo-EM density of UL32, where one pentamer is shown in green (147,337 particles, C1). (**b**) Views of the UL32 pentamer model as ribbon helices, with each protomer shown in a different shade of green. Center and right views show the pentamer model inside the UL32 tripentamer map blurred to B-factor 2000 and shown as a transparent surface. (**c**) UL32 protomer colored by the Cα RMSD to ORF68 (PDB ID: 6XF9), blue is 1 Å, white is 3 Å, red is 5 Å, and yellow are residues without RMSD values where there are insertions relative to ORF68. (**d**) Views of the UL32 tripentamer model as tube helices, with the three pentamers shown in medium green, light green, and grey. Top row shows the rotation around the 3-fold symmetry axis, while the bottom row shows views from along the 3-fold symmetry axis and a view along the local 2-fold symmetry axis at the tripentamer interface.

The tripentamer consists of three pentameric rings arranged around a 3-fold symmetry axis (**Fig 3d**). The interface between two pentameric rings involves four protomers (two protomers from each pentamer) engaged in a local 2-fold symmetric interaction. The tripentamer architecture lacks global 2-fold symmetry because the three pentamers are splayed outward by ∼18° relative to a regular prism. This arrangement forms a triangular frustum of interior space, where the smaller equilateral triangle has a length of ∼70 Å, and the larger widens to ∼100Å. For the purposes of our discussion, we define the wider opening as the “top” of the tripentamer.

At the top of the tripentamer, one protomer from each pentameric ring does not contact neighboring pentamers (**Fig 3d**). In the UL32 consensus map, one of the three top protomers had weaker density and lower local resolution (**Fig S5e**). Further 3D classification revealed that the majority of UL32 particles form complete tripentamers (15-mers, ∼60%), while the remainder form incomplete tripentamers containing two 5-membered rings and an incomplete ring missing one or more protomers (**Fig S6a**). Over 40% of incomplete tripentameric particles (17% of total particles) formed a 14-mer, with a twisted, 4-membered incomplete ring. This UL32 4-membered incomplete ring is reminiscent of the UL52 4-membered incomplete ring, but close inspection shows that while the UL52 4-mer twist has a consistent rise, the UL32 4-mer twist is asymmetric, arising largely from the rotation of one protomer (**Figs S6b, S6c**). As in UL52, the twist arises from hinging at the loop-pocket motif at the protomer interface (**Fig S6d**).

Since the UL32 cryo-EM sample was prepared in the presence of 30 bp dsDNA, we carried out further classification to resolve UL32 interactions with dsDNA. Symmetry expansion followed by 3D classification with a mask focused on the potential dsDNA region revealed two classes of particular interest (**Fig S6e**). In one class (30% of all particles), a rod-shaped extra density spans the open interior space of the tripentamer between the central channels of two pentamers. A 30 bp dsDNA model, displayed as a molecular surface at 10 Å resolution, approximates the length and shape of this additional density; the absence of clear helical character in this density suggests that the potential dsDNA density is rotationally averaged (**Fig S6f**). In another class, the extra density near the central channel has a distinct right-handed twist that fits well with the minor groove of a B-form DNA model (**Fig S6g**). In both classes, the extra density inserts into, but does not entirely span, the central channel of the UL32 pentameric ring.

### Residues at the tripentamer interfaces contribute to infectious virion production in HSV-1

The tripentameric architecture creates a distinct interface between the protomers of adjacent pentameric rings. Each of the three locally 2-fold symmetric inter-pentamer interfaces has two contact points between the same α-helix from protomers of adjacent pentamers (residues 523-536) (**Fig 4a**). Two copies of residue C535 are separated by ∼8 Å (Cα-Cα distance) with connecting map density visible at low to moderate contour levels (**Figs 4a**, **S7a**). A lysine residue (K532) on this helix could contribute to stabilization of the tripentamer in our crosslinked sample, as EGS targets primary amines^30^. HCMV UL52 and KSHV ORF68 have an α-helix in the equivalent position to the UL32 tripentamer interface (UL32 residues 523-536, **Fig S7b**). An equivalent cysteine to UL32 C535 is conserved in nearly all α/β-herpesvirus homologs but is absent in the γ-herpesvirus homologs (**Fig S1d**). Nearby in the tripentamer interface, a segment of non-continuous protein density can be modeled as a short α-helix from the N-terminus of UL32 (residues 5-11) that is sufficiently long to reach from the last modeled residue (P19) to the tripentamer interface (**Fig S7a**). While a subset of human herpesvirus packaging accessory factors have N-terminal extensions, they are poorly conserved, with the UL32 N-terminal extension sharing sequence similarity only with that of the closely related α-herpesvirus HSV-2

**Fig 4.**
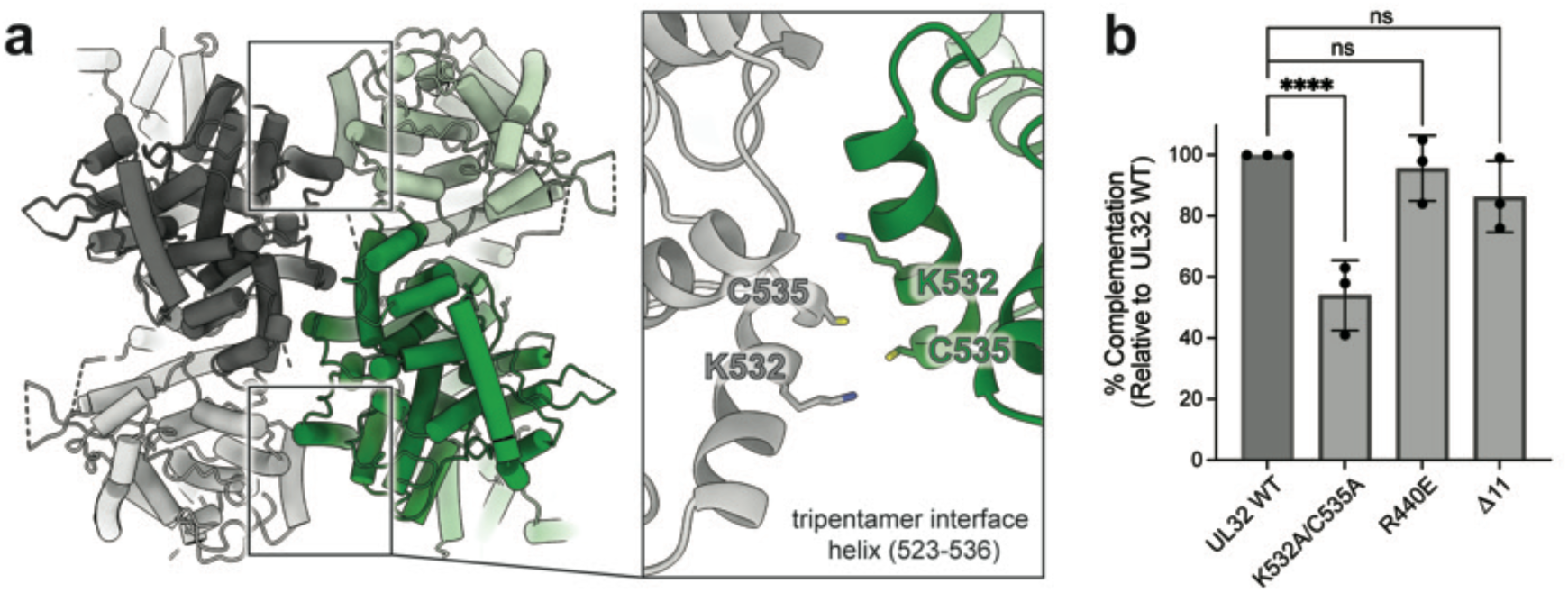
The tripentamer interface contributes to production of infectious virions in HSV-1. (a) Tripentamer interface viewed along the local 2-fold symmetric interaction axis. Inset shows model details. (b) HSV-1 transient complementation assay of UL32 tripentamer interface mutants from three independent biological replicates. Complementation efficiency was analyzed using a one-way ANOVA followed by Dunnett’s multiple comparisons test. Error bars represent SD. \*\*\*\**P* < 0.0001, ns, not significant.

(**Fig S1e**).

We wanted to understand if residues at the tripentamer interface are required for UL32 function in HSV-1 infection. Loss of UL32 prevents viral genome packaging and infectious virion production^20^. We used a previously described transient complementation assay^21^ to test the effect of mutations at the UL32 tripentamer interface (**Fig S7c**). Loss of the N-terminal α-helix (Δ11) or a charge swap mutation at the interface (R440E) (**Figs S7a, S7d**) had minimal effect on infectious virion production or UL32 expression (**Figs 4b**, **S7e, S7f**). However, mutation of K532A/C535A reduced infectious virion production by half (**Fig 4b**), suggesting that these residues play a subtle but reproducible role in the viral life cycle.

### All herpesvirus packaging accessory factors contain a conserved core fold stabilized by zinc fingers

The structure of ORF68, the KSHV packaging accessory factor, revealed that each protomer contained three CCCH-type zinc fingers (ZnF)^28^. The first two zinc fingers are present in our structures of HSV-1 UL32 and HCMV UL52, with identical coordination and highly similar local structures conserved across all homologs (**Fig 5a**). ZnF1 forms part of the pocket of the pocket-loop motif of the protomer interface. Unlike the α- and γ-herpesvirus accessory factors, HCMV UL52 lacks a third zinc finger (**Fig 5b**), consistent with sequence alignments indicating that this motif is absent in the β-herpesviruses^28^. In the position where ORF68 and UL32 have ZnF3, UL52 has two residues (C653, H333) that could coordinate zinc, but it lacks two additional residues to fulfill tetrahedral coordination (**Figs 5b**, **S1d, S1f**). The third zinc finger in ORF68 consists of residues C415, H452, and the unusual use of adjacent cysteines C191 and C192 (**Fig 5b**). These four metal-binding residues are not conserved across herpesvirus homologs (**Figs S1d, S1f**), but two adjacent cysteines were identified in HSV-1 UL32 (C308 and C309) and proposed to be the third and fourth metal-binding residues^28^. Our structure shows that while C308 does bind Zn3, C309 is not involved in coordination (**Fig 5b**). Instead, C297 – present on the same loop as C308 – is the fourth coordinating residue of ZnF3 (**Figs 5b**, **S5h**).

**Fig 5.**
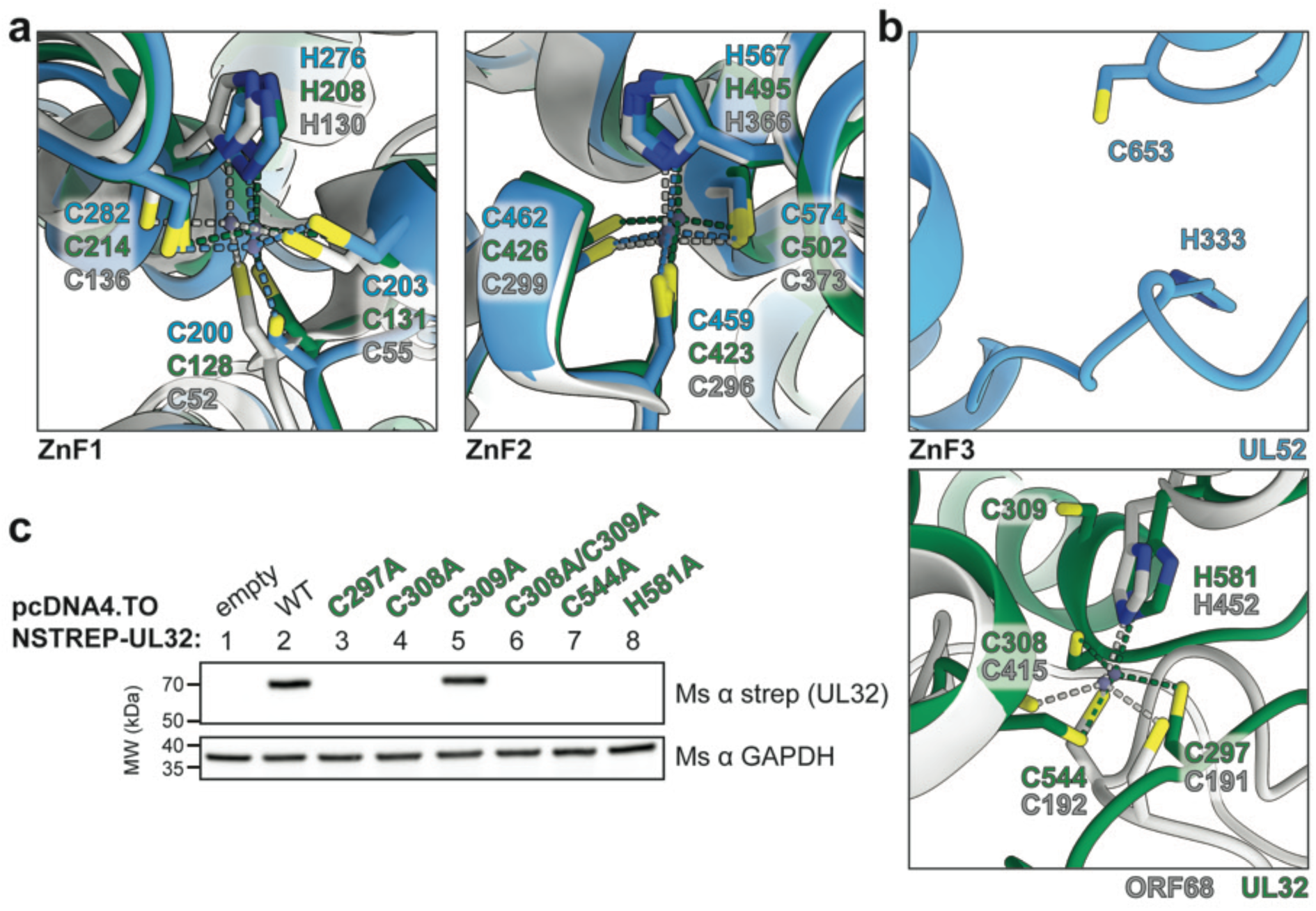
**The herpesvirus packaging accessory factor contains a conserved core fold stabilized by zinc fingers.** (**a-b**) Zinc finger (ZnF) coordination in UL52 (blue), UL32 (green) and ORF68 (grey). Zn^2+^ ions are shown as blue-grey spheres connected by dashed lines to coordinating cystine and histidine residues, shown as sticks. (**c**) Western blot for expression of UL32 WT and ZnF3 mutants by transient transfection, representative of three independent biological replicates.

The zinc finger motifs in ORF68 and UL32 were previously found to be required for the structural stability of these proteins *in vivo*, and mutations in ZnF1 (C128A), ZnF2 (C502A) or ZnF3 (C544A, H581A, and the double mutant C308A/C309A) greatly reduced UL32 levels in transfected HEK293T cells^28^. We show that mutation to alanine of the four metal-binding residues of ZnF3 observed in our structure (UL32 C297, C308, C544, and H581) reduced UL32 expression, while C309A alone had no effect (**Fig 5c**). These data support our structural identification of the residues comprising the essential zinc fingers required for stability of packaging accessory factors across the *Herpesviridae*.

### A conserved positively charged central channel is required across the Herpesviridae

Our structures show that packaging accessory factors from all three herpesvirus subfamilies can assemble into rings with pentameric symmetry, creating a channel in the center of the ring. Each protomer contributes a well-structured α-helix and a semi-structured loop that form the surface of the central channel (**Fig 6a**). These central channel regions are studded with relatively well-conserved lysine and arginine residues, creating a positively charged, structurally conserved channel (**Figs 6a**, **S1d, S1f**). The central channel in ORF68 was previously shown to be essential for viral genome packaging, as mutation of positive charges in the central channel (the single mutation K435A and triple mutation K174A/R179A/K182A) phenocopied total loss of ORF68^28^.

**Fig 6.**
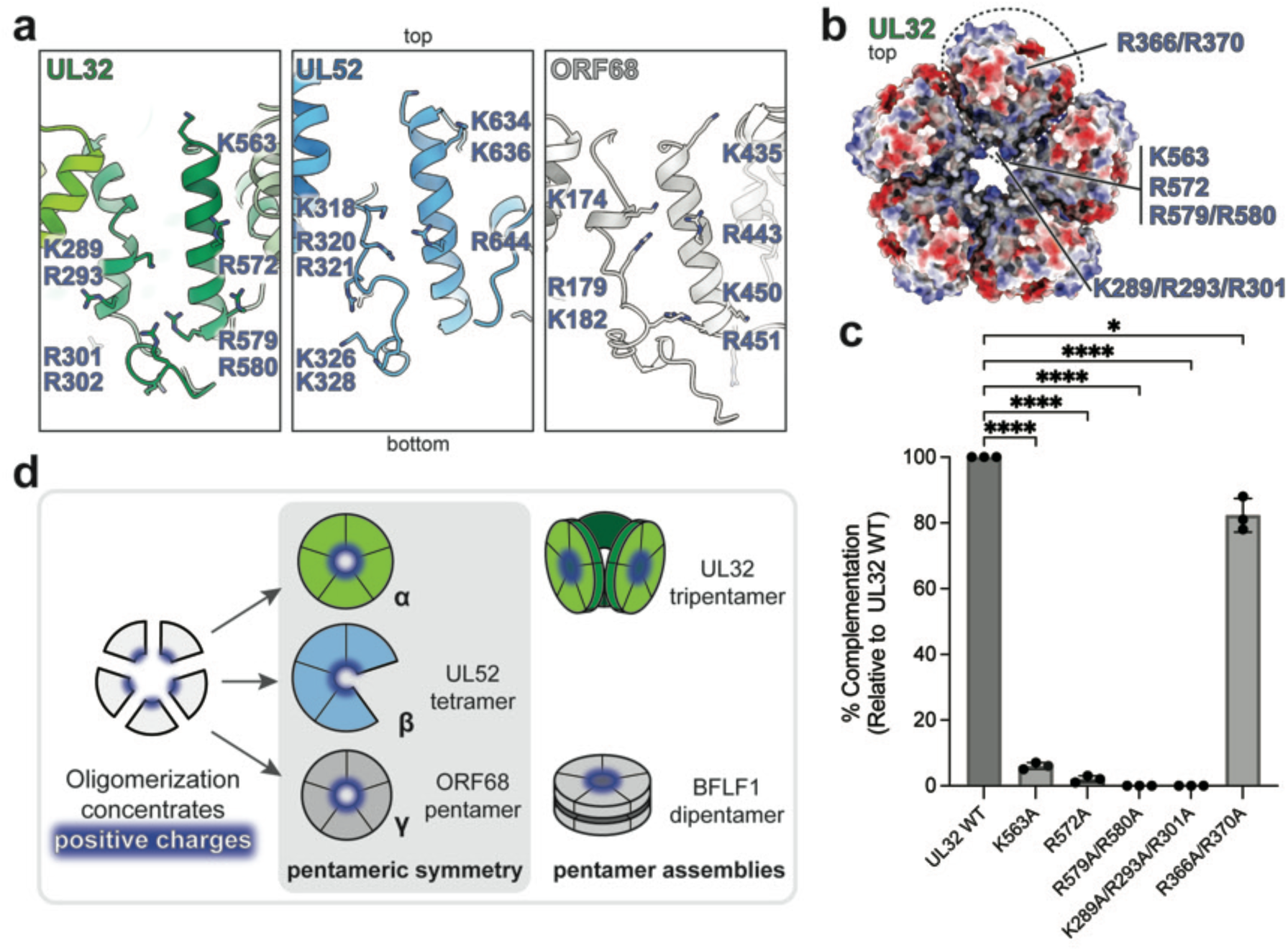
**A positively charged central channel is conserved and required across the *Herpesviridae.*** (**a**) View of central channel of UL32, UL52, and ORF68, shown as cartoon with lysine and arginine residues displayed as sticks for one protomer. (**b**) Location of UL32 mutations. UL32 pentamer model shown as electrostatic surface (Coulombic potential) viewed from the top of the ring, with a dashed line encircling one protomer. (**c**) HSV-1 transient complementation assay of UL32 central channel mutants and control mutant from three independent biological replicates. Complementation efficiency was analyzed using a one-way ANOVA followed by Dunnett’s multiple comparisons test. Error bars represent SD. \*\*\*\**P* < 0.0001, \**P* < 0.05. (**d**) Model of assembly of herpesvirus packaging accessory factors, where assembly serves to concentrate positive charge within a central channel.

We tested if the positively charged central channel residues of UL32 were required to produce infectious virion in HSV-1 using the transient complementation assay (**Fig S7c**)^21^. We generated the UL32 mutants K563A (positionally equivalent to ORF68 K435, at the top of the channel), R572A (in the middle of the channel), R579A/R580A (at the bottom of the channel), and K289A/R293A/R301A (extending throughout the central channel, positionally equivalent to ORF68 K174A/R179A/K182A) (**Fig 6a**, **6b**). All mutations drastically decreased production of infectious virus particles without affecting UL32 expression (**Figs 6c**, **S7e, S7f**). Mutating a single positively charged residue (K563A or R572A) reduced viral yield by an order of magnitude, and the double- and triple-point mutants were equivalent to total loss of UL32 (**Figs 6c**, **S7f**). This effect was specific to mutation of positively charged residues in the central channel, as mutation of two positively charged residues on the outer surface of UL32 (R366A/R370A) did not significantly impact virus production (**Figs 6b, 6c**, **S7e, S7f**). Thus, UL32 requires a positively charged central channel, and the structural conservation of this feature is suggestive of a shared function across the *Herpesviridae* (**Fig 6d**).

## DISCUSSION

Our structures of packaging accessory factor homologs from across the herpesvirus family illuminate the conserved architecture of this essential viral factor. The herpesvirus packaging accessory factor homologs share a core fold whose surface is punctuated with insertions and variations. Some sites of difference are common to α/β-herpesvirus homologs, while others are subfamily-specific (**Fig S1a**). For example, both UL32 and UL52 have an additional α-helix on the top peripheral surface of the ring compared to ORF68, as well as an N-terminal extension and two disordered loops on the top surface. β-herpesvirus homologs furthermore have an additional loop on the side of the ring, while α-herpesvirus homologs have an additional disordered loop and a shifted α-helix on the bottom of the ring. The variability of the outer surface of the ring suggests that these regions are involved in virus-specific interactions. Indeed, factors from different subfamilies are not interchangeable: HSV-1 UL32 and HCMV UL52 cannot complement for loss of ORF68 during KSHV infection^28^. Even homologs from within the γ-herpesvirus subfamily (EBV BFLF1, MHV68 muORF68) are unable to fully replace ORF68 in KSHV^28^.

Despite a common core fold, the recombinantly purified packaging accessory factor homologs displayed a difference in their propensity to form stable assemblies *in vitro* despite similar purification and buffer conditions. Regardless of the stability of this assembly, all of the homologs we have examined have the ability to form oligomers with pentameric symmetry. The γ-herpesvirus homologs ORF68 and BFLF1 form stable pentamers^28^. In contrast, UL52 prefers to exist as an incomplete ring with pentamer-like symmetry, while UL32 can assemble into pentameric rings. A structurally conserved loop-pocket motif acts as a hinge to modify the orientation of interacting protomers, but the degree of rotation is constricted such that the ring ultimately maintains near-5-fold symmetry. The differences in quaternary structures of packaging accessory factors across the herpesvirus subfamilies may be less pronounced during infection, as the relative stability of the rings may be modulated through interactions with other viral factors (e.g., terminase or capsid) or biophysical conditions within replication compartments.

Although the packaging accessory factor homologs have poor surface conservation overall and differ in the propensity to form stable oligomeric rings, they likely serve the same function in viral genome packaging given that viruses lacking these factors have identical phenotypes^20,22–24^. We previously proposed potential roles for the herpesvirus packaging accessory factor, speculating that it may act as a scaffold for assembly of the terminase packaging motor, aid association between the portal and terminase, or coordinate other essential pieces of the packaging or late gene transcription machinery^28^. While the precise mechanistic contribution of the packaging accessory factor remains unknown, our structural data on UL32 and UL52 suggests that the shared ability to form a pentameric ring is tied to its essential and conserved function. Indeed, ring formation concentrates positive charges within a central channel, and these positively charged residues are critical for successful herpesvirus packaging.

Could this central channel be used to bind the viral dsDNA genome? While the central channel, 25 Å at its narrowest point, could technically accommodate dsDNA, it is unlikely to allow an intact ring to slide along dsDNA. In contrast, the sliding clamp involved in DNA replication (i.e., proliferating cell nuclear antigen (PCNA)) forms a symmetric ring with a positively charged central channel ∼35 Å in diameter, easily accommodating duplex DNA^31,32^. In an interesting analogy, herpesviruses encode their own sliding clamp processivity factor (HSV-1 UL42/HCMV UL44/KSHV ORF59) that shares a common PCNA-like fold but whose oligomeric state varies across the herpesviruses and forms an incomplete ring^33–36^. A surprising finding in our study was the novel tripentameric quaternary assembly of UL32. Interestingly, ORF68 has also been shown to form discrete higher molecular weight bands in EMSA on dsDNA pieces 30 bp and longer^23,28^, suggestive of supra-pentameric assemblies in other herpesviruses. We speculate that the formation of higher-order assemblies could play a role in viral genome packaging, and future work to uncover the precise role of the packaging accessory factor should illuminate the mechanistic contribution of distinct oligomeric states.

## MATERIALS AND METHODS

### Plasmids

ORF68 and UL32 with N-terminal Twin-Strep tags and HRV 3C protease cleavage sites (“TSP”) were subcloned from their respective pUE1-TSP plasmids (Addgene #162650, 162658) into pLIB (Addgene #80610) digested with BamHI/HindIII to generate pLIB-TSP-ORF68 and pLIB-TSP-UL32 (Addgene #250565, 250566). The coding region of UL52 was subcloned from a pcDNA4/TO-2xStrep vector (Addgene #162628) into a pHEK293 UltraExpression I vector (Clontech) that encodes an N-terminal Twin-Strep tag and HRV 3C protease cleavage site (pUE1-TSP) to generate pUE1-TSP-UL52. Then, TSP-UL52 was subcloned into pLIB as described above to generate pLIB-TSP-UL52 (Addgene #250567). UL32 mutants used for transient expression experiments (Addgene 250568-250578) were generated by inverse PCR of pcDNA4/TO-2xStrep-UL32 (Addgene #162629). Plasmids expressing UL32 C308A/C309A, C544A, and H581A were previously described (Addgene #162646-162648).

### Expression and purification of recombinant ORF68, UL52, UL32

TSP-ORF68 and TSP-UL52 were expressed in Sf9 insect cells using the Bac-to-Bac Baculovirus Expression System (Thermo Fisher Scientific). pLIB-TSP-ORF68 or UL52 was transformed into *E. coli* DH10Bac competent cells, from which bacmid was isolated and transfected into Sf9 cells with Cellfectin II (Thermo Fisher Scientific) to generate baculovirus. Sf9 cells were infected with baculovirus for protein expression and purification. Three days later, cells were pelleted at 1000 × g for 5 min at 4°C and resuspended in 30 mL lysis buffer (100 mM Tris-HCl pH 8.0, 300 mM NaCl, 5% glycerol, 0.5% CHAPS, 1 μg/mL avidin, 1 mM dithiothreitol (DTT), cOmplete™ EDTA-free Protease Inhibitor Cocktail (Millipore Sigma)) for 30 min at 4°C. The cell suspension was sonicated and the lysate clarified by ultracentrifugation at 50,000 × g for 30 min. The lysate was filtered through a 0.45 μm filter before purification by a gravity column with Strep-Tactin XT 4Flow resin (IBA Lifesciences). The column was washed with wash buffer (100 mM Tris pH 8.0, 300 mM NaCl, 0.1% CHAPS, 1 mM DTT), and the protein was eluted in wash buffer containing 50 mM biotin. The protein was concentrated using a 30 K Amicon Ultra-15 concentrator (Millipore) before injection onto a Superose 6 Increase 10/300 GL (24 mL, 5kDa-5000KDa, 500 μL load) equilibrated in SEC buffer (20 mM HEPES pH 7.6, 100 mM NaCl, 5% glycerol, 1 mM DTT). The peak fraction was pooled, concentrated, and exchanged into running buffer containing 10% glycerol for freezing using a 30 kDa MWCO Amicon Ultra-15 centrifugal filter (Millipore).

TSP-UL32 was expressed as above, with the exception that Strep-Tactin Sepharose resin was used for purification and the elution buffer contained 400 mM NaCl and 2.5 mM desthiobiotin. Additionally, the SEC buffer contained 500 mM NaCl. Protein used in SEC-MALS experiments were purified as above, but recombinant expression occurred in High Five insect cells for 52 h (Thermo Fisher Scientific), lysis buffer contained 20 μg/mL DNase I (Gold Biotechnology), buffers contained 1 mM Tris(2-carboxyethyl)phosphine hydrochloride (TCEP) instead of DTT, and proteins were eluted from the Strep-Tactin column by cleavage with HRV-3C protease.

### SEC-MALS

The oligomeric state of purified packaging accessory factors was determined using Size Exclusion Chromatography (SEC) coupled with Multi-Angle Light Scattering (MALS). SEC was performed with a Superdex 200 Increase column. Each sample injection consisted of 100 μL of 0.5-1 mg/mL purified protein in running buffer (50 mM HEPES pH 7.6, 100 mM NaCl, 5% glycerol, 1 mM TCEP for UL52 and ORF68; for UL32 the NaCl concentration was 500 mM) at a flow speed of 0.4 mL/min. MALS was monitored with a DAWN Heleos II light scattering detector and an Optilab T-rEX refractive index detector (both from Wyatt Technology). Light scattering data were collected at 1 second interval and analyzed with ASTRA6 software (Wyatt Technology).

### Electrophoretic mobility shift assay

Fluorescein-labeled dsDNA probes (Integrated DNA Technologies, previously described in^28^, except for the 67 bp probe: /56-FAM/cgccgccgggcctgcggcgcctcccgcccggg Catggggccgcgcgccgcctcagggcccggcgcgg + ccgcgccgggccc tgaggcggcgcgcggccccatgcccgggcgggaggcgccgcaggcccggcggcg), were prepared as 2x stocks (20 nM) in binding buffer (100 mM NaCl, 20 mM HEPES pH 7.5, 5% glycerol, 0.05% CHAPS, 1 mM DTT). UL32 was diluted in binding buffer containing 0.2 mg/mL Bovine Serum Albumin (BSA). Binding reactions were prepared by mixing equal volumes of probe DNA and protein solutions, resulting in final concentrations of 10 nM DNA probe, 100 mM NaCl, 20 mM HEPES pH 7.5, 5% glycerol, 0.05% CHAPS, 0.1 mg/mL BSA, and variable concentrations of UL32. Concentrations are indicated as calculated for a UL32 protomer. Samples were incubated at room temperature for 30 min prior to electrophoresis on a 5% polyacrylamide (29:1 acrylamide:bis-acrylamide)/ 1x Tris borate gel at 2W at 4°C. Gels were imaged using an Amersham Typhoon imaging system (Cytiva).

### Crosslinking of UL32 and negative stain EM

UL32 (4 µM) was incubated with 0.5-4 mM ethylene glycol bis(succinimidyl succinate) (EGS) for 30 min on ice. For samples with DNA, UL32 (4 µM) was incubated with 5 µM dsDNA probe for 30 min at room temperature prior to crosslinking. DNA probes were the same as described for EMSA. Crosslinking products were resolved and visualized by Coomassie stained 4-15% TGX SDS-PAGE (BioRad) and by negative stain microscopy. Samples for negative staining were diluted to a concentration of 250 nM, then applied to glow-discharged, continuous carbon EM grids (Electron Microscopy Science (EMS), CF400-Cu-50) and stained with 2% uranyl acetate (EMS, 22400-2). EM grids were imaged in a Talos L120C transmission electron microscope operated at 120 kV. Micrographs were collected manually at magnifications of 36,000×, 45,000x, or 73,000x

### Cryo-EM sample preparation and data collection

Cryo-EM samples were prepared with purified UL52 (7 µM final concentration) and purified UL32 (15 µM final concentration). Prior to freezing UL32, 4 µM UL32 was incubated with 5 µM 30 bp dsDNA (as used for EMSA) for 30 min at room temperature, chemically crosslinked with 2 mM EGS for 30 min on ice, quenched with 50 mM Tris-HCl, and concentrated to 15 µM in a 30 kDa MWCO Amicon Ultra-15 centrifugal filter.

Samples (4 µL) were applied to C-flat R2/1, 300 mesh, holey carbon copper grids (Electron Microscopy Sciences) after glow discharging the grids at 11 mA for 30 sec (Pelco). CHAPSO detergent was added to a final concentration of 0.05% just before grid freezing to ensure uniform ice thickness and particle distribution. Samples were vitrified with a Vitrobot Mark IV system (Thermo Fisher Scientific) maintained at 10°C and 100% humidity. Grids were blotted at blot force 1 for 6 sec and plunge frozen in liquid ethane.

Grids were screened for optimal ice thickness and particle concentration. Data collection was carried out from a single screened grid using a 300 kV Titan Krios cryo-transmission electron microscope (Thermo Fisher Scientific) equipped with a K3 camera (Gatan) and an imaging energy filter (Gatan) operated at a slit width of 15 eV. The dataset was collected in counting super-resolution mode with a nominal magnification of 81,000x leading to a physical pixel size of 1.07 Å (super-resolution pixel size is 0.535 Å). The data were collected at a dose rate of 15 e-/pixel/sec with a total electron dose of 50 e-/Å2 applied over 40 frames and a targeted defocus range of −1.0 µm to −2.0 µm.

### Cryo-EM data processing and model building

The cryo-EM data processing workflow was carried out in CryoSPARC v4^37^. Movie frames were motion corrected using Patch motion correction and binned two-fold to yield a motion-corrected micrograph stack with a pixel size of 1.07 Å. Micrographs were manually curated and an initial round of particle picking was performed using blob picker on a subset of 1,000 micrographs. A 2D classification job on the initial particle stack yielded 2D templates that were used for template-based particle picking on the entire dataset. The template-picked particle stack was curated using 2D classification and used for *ab initio* reconstruction and heterogenous refinement of the partial stack. For UL52, classes for the 3-mer and 4-mer were separately refined using additional 2D classification and heterogenous refinement. No symmetry restrains were applied to maps (C1) though in the case of UL32 C3 symmetry could be applied and increased the resolution of the map (C1 3.09 Å vs. C3 2.89 Å). Final particle stacks were polished using Reference-based motion correction and used for NU-refinement to create the final reconstructions. Final reconstructions were uploaded to the Electron Microscopy Data Bank (EMDB).

Further analysis of the UL32 consensus particle stack to resolve incomplete tripentamers (i.e., 14- vs. 15-mers) was done with iterative 3D classification with focus masks, homogenous, and non-uniform refinement. The UL32 consensus particle stack was further analyzed for the presence of dsDNA probe included in the cryo-EM sample. Classes with additional density consistent with the DNA were generated by C3 symmetry expansion followed by iterative 3D classification with a focus mask and homogenous refinement.

Model building was done in Coot^38,39^ using starting templates generated with AlphaFold3^40^. After initial rigid-body docking, residues were manually fit into the high-resolution map using real-space refinement modules. Rounds of real-space refinement in PHENIX^41^ were carried out and the outliers were manually corrected in Coot before deposition in the Protein Data Bank (PDB).

### Structure analysis and sequence alignment

Structures were visualized and figures generated with ChimeraX^42^. Electrostatic surfaces are shown with coulombic coloring. Structural analysis included two additional models generated by Alphafold3^40^: in Fig 6a an ORF68 model was used to represent regions that were unmodeled in the ORF68 crystal structure (6XF9); and in S6f, S6g Fig a 30 bp dsDNA model was docked into densities that could likely be attributed to dsDNA. The sequence of packaging accessory factors from the nine human herpesviruses were obtained from UniProt^43^ (HSV-1, P10216; HSV-2, P89455; VZV, P09282; HCMV, P16793; HHV-6A, P52463; HHV-6B, Q9QJ33; HHV-7, P52464; EBV, P03184; KSHV, F5HF47) and aligned using Clustal Omega.

### Cell lines and viruses

HEK293T cells (ATCC CRL-3216) were maintained at 37°C with 5% CO_2_ in Dulbecco’s Modified Eagle Medium (DMEM) supplemented with 10% fetal bovine serum (FBS) (LifeTech). Vero cells (ATCC CCL-81) and the UL32-complementing cell line 158^20^ were maintained at 37°C with 5% CO_2_ in DMEM supplemented with 5% FBS and 1% penicillin-streptomycin. The HSV-1 UL32-null mutant virus *hr*64FS was previously described^21^.

### Transient transfection and western blot analysis

HEK293T cells were plated in 6-well plates and transfected after 24 hours at 70% confluency with polyethylenimine (PEI). Cells were harvested 24 hours later by washing in DPBS, then resuspending cells in Dulbecco’s Phosphate-Buffered Saline (DPBS) prior to centrifugation at 500 x g for 5 minutes at 4°C. Cell pellets were stored at −80°C. Samples were prepared for western blot analysis by resuspension in lysis buffer (150 mM NaCl, 50 mM Tris pH 8.0, 1 mM EDTA, 0.5% NP-40, cOmplete protease inhibitor [Roche]) and rotating at 4°C for 1 hour. Insoluble material was removed through centrifugation at 15,000 x g for 10 minutes at 4°C, then the total protein concentration in the cell lysate was determined by a Bradford protein assay. Lysate was mixed with Laemelli loading buffer and 30 μg of total protein was separated on a 4-15% TGX SDS-PAGE gel (Bio-Rad) prior to transfer onto PVDF membrane. Membranes were blocked with 5% milk in TBST buffer (Tris-buffered saline, 0.2% Tween 20) and incubated overnight at 4°C with mouse monoclonal anti-GAPDH (1:5000, cat. no. AM4300 clone 6C5, ThermoFisher) or mouse monoclonal anti-Strep Tag II (1:500, cat. no. PIMA537747, Invitrogen) primary antibodies. The next day, the membrane was washed in TBST and incubated for 1 hour at room temperature with secondary antibody (polyclonal goat anti-mouse HRP, 1:5000, Southern Biotech). The blot was washed with TBST then incubated with Clarity Western ECl substrate (Bio-Rad) and imaged on an Azure Biosystems 600 imager.

### Transient complementation assay and western blot analysis

The UL32 transient complementation assay using *hr*64FS was performed as previously described^21^. Briefly, Vero cells grown to ∼75% confluence in a 12-well plate were transfected with 0.5 μg of empty, wild-type, or mutant plasmid using the Lipofectamine 2000 reagent (Thermo Fisher) according to the manufacturer’s protocol. At 14-16 h posttransfection, cells were superinfected with *hr*64FS at an MOI of 5 PFU. At 24 h post infection, media and cells were harvested, centrifuged to separate virus-containing supernatant from cells. Cell pellets were washed with 1x PBS, reconstituted in 1x SDS loading buffer and subjected to western blot analysis for protein expression as described above. A plaque assay to quantify total viral yield was performed by infecting 158 (UL32-complementing) cells with serial dilutions of the viral supernatants. Infected cells were overlayed with 2% w/v methylcellulose prepared in 2% FBS DMEM and incubated for 72 h. Cells were fixed with final concentration of 2% formaldehyde and stained with 1% crystal violet solution. Plaques were counted and viral titers were determined. The percent complementation was calculated by dividing the titer obtained for the mutant plasmid by the titer obtained for the wild-type plasmid and multiplying by 100. The background titers from the empty plasmid samples were subtracted. The following primary antibody were used for western blot analysis: rabbit polyclonal anti-UL32 (1:500; antibody to synthetic antigenic peptide^21^ and mouse monoclonal anti-gamma tubulin (1:5,000, Sigma, T6557) used as a loading control.

## ACKNOWLEDGEMENTS

We are thankful to members of the Didychuk lab and Xiong lab for feedback. Negative stain electron microscopy and cryo-EM screening were conducted at the Yale Cryo-EM Resource, which is funded in part by an NIH grant (S10OD023603) awarded to F. Sigworth. Cryo-EM data were collected at The Laboratory for BioMolecular Structure (LBMS), which is supported by the DOE Office of Biological and Environmental Research (KP1607011). Molecular graphics and analyses performed with UCSF ChimeraX, developed by the Resource for Biocomputing, Visualization, and Informatics at the University of California, San Francisco, with support from National Institutes of Health R01-GM129325 and the Office of Cyber Infrastructure and Computational Biology, National Institute of Allergy and Infectious Diseases.

## DATA AVAILABILITY

Coordinates and cryo-EM maps have been deposited into the Protein Data Bank and Electron Microscopy Data Bank under the following accession codes: HCMV UL52 4-mer (PDB 9ZM2 [doi.org/10.2210/pdb9ZM2/pdb], EMD-74419 [ebi.ac.uk/emdb/EMD-74419]), HCMV UL52 3-mer (EMD-74418 [ebi.ac.uk/emdb/EMD-74418]), HSV-1 UL32 tripentamer (PDB 9ZLY [doi.org/10.2210/pdb9ZLY/pdb], EMD-74415 [ebi.ac.uk/emdb/EMD-74415]). The previously published model of ORF68 used for comparative structural analysis is available in the Protein Data Bank using accession code PDB 6XF9 [doi.org/10.2210/pdb6XF9/pdb].

## FUNDING

This work was supported by an NIH DP2 AI171113 award and Damon Runyon Cancer Research Foundation Dale F. Frey Award for Breakthrough Scientists awarded to A.L.D, with partial salary support to A.L.D. from both grants. E.J.B. received salary support from an American Cancer Society Postdoctoral Fellowship (PF-24-1322561-01-RMC). L.M.M. received salary support from a Yale Endowed Postdoctoral Fellowship and a Deutsche Forschungsgemeinschaft Walter Benjamin Programme Fellowship (561072518). The funders had no role in study design, data collection and analysis, decision to publish, or preparation of the manuscript.

## COMPETING INTERESTS

The authors declare that they have no conflict of interest.

## SUPPLEMENTARY FIGURE LEGENDS

**S1 Fig.**
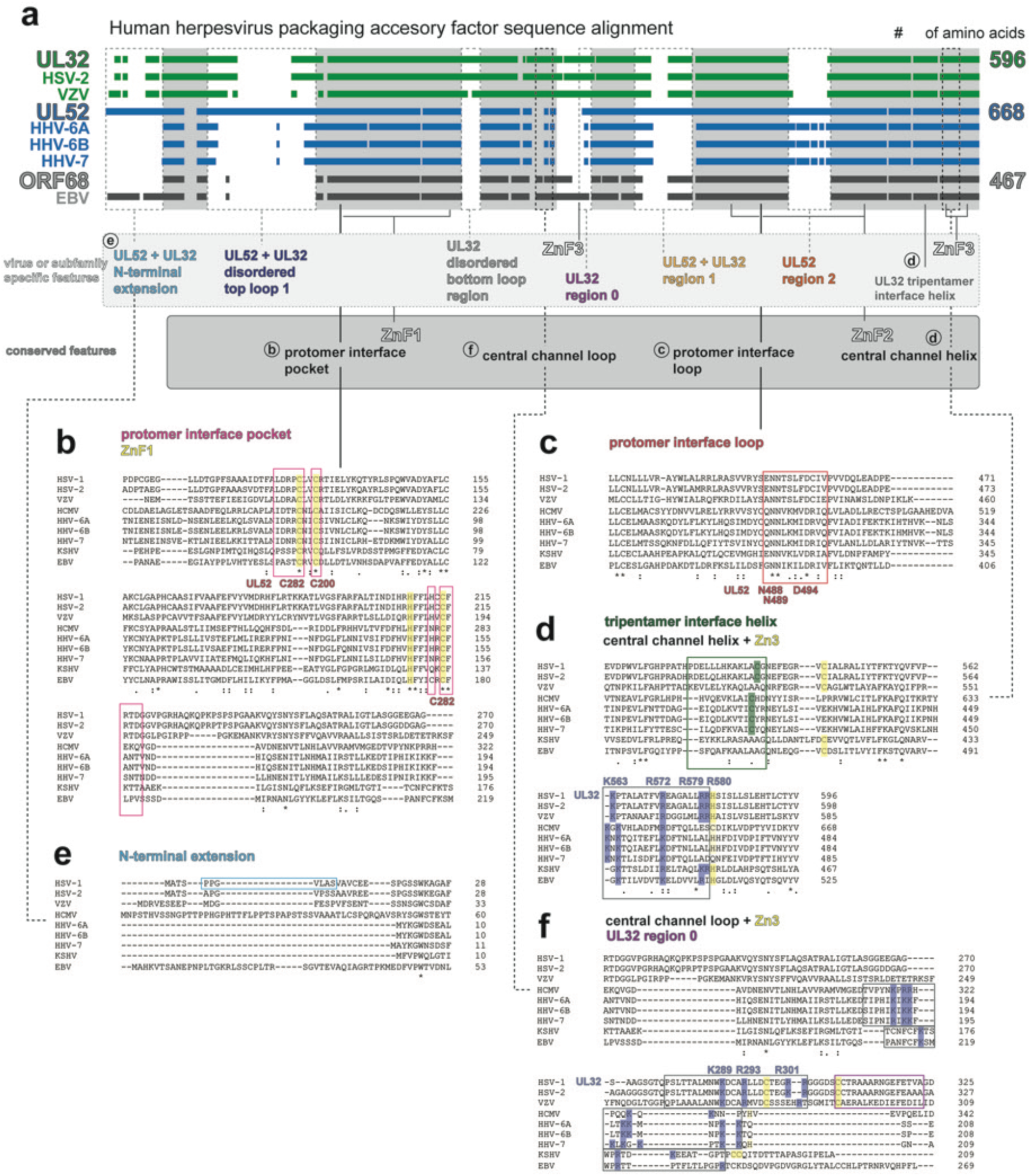
Sequence alignment of packaging accessory factors for the nine human herpesviruses. (**a**) Schematic overview of sequence alignment for packaging accessory factors from the nine human herpesviruses shows conserved features and subfamily or virus specific features. (**b-f**) Details of sequence alignment for conserved and divergent features in the α-, β-, and γ-herpesviruses with relevant features boxed and highlighted. In **b** and **c**, the universally conserved residues for the protomer interface loop are labeled for HCMV UL52. In **d**, **f** conserved positively charged residues in the central channel are highlighted in blue for all homologs and labeled for the HSV-1 UL32 residues that this study shows to be functionally important.

**S2 Fig.**
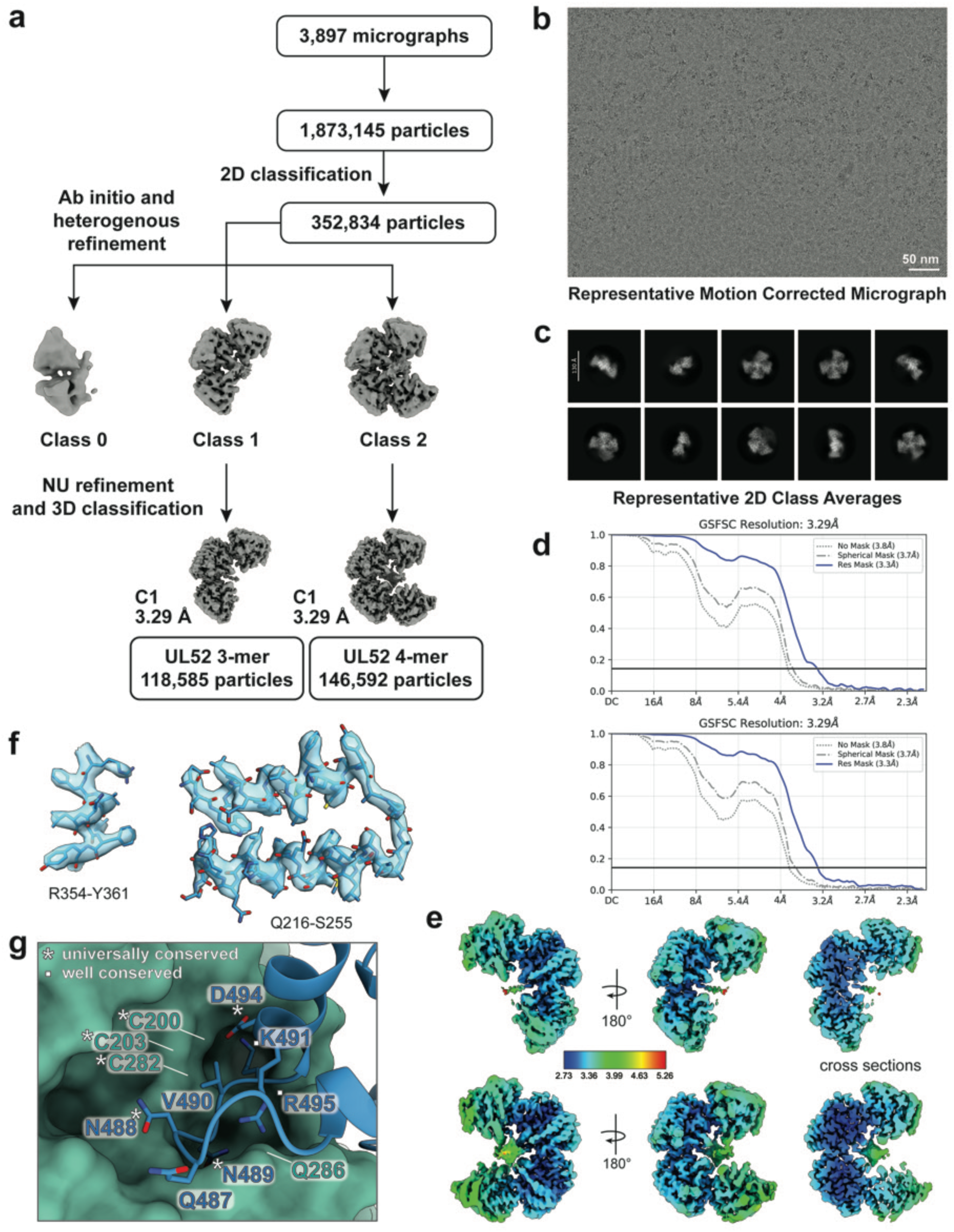
Cryo-EM data analysis for HCMV UL52. (**a**) Overall data processing workflow for the HCMV UL52 cryo-EM reconstruction. (**b-c**) Representative micrograph with scale bar (50 nm) and 2D classes. (**d**) Gold-standard Fourier Shall Correlation (GS-FSC) curve for the UL52 3-mer (top) and 4-mer (bottom). (**e**) Local resolution for the 3-mer and 4-mer reconstructions. (**f**) Representative examples for the cryo-EM map and the corresponding modeled regions. Map has transparent surface; model is shown with sticks and heteroatom coloring (red for oxygen, yellow for sulfur, blue for nitrogen). (**g**) Residues in the protomer interface, colored as in **Fig 2e** but with loop residues shown as sticks with heteroatom coloring.

**S3 Fig.**
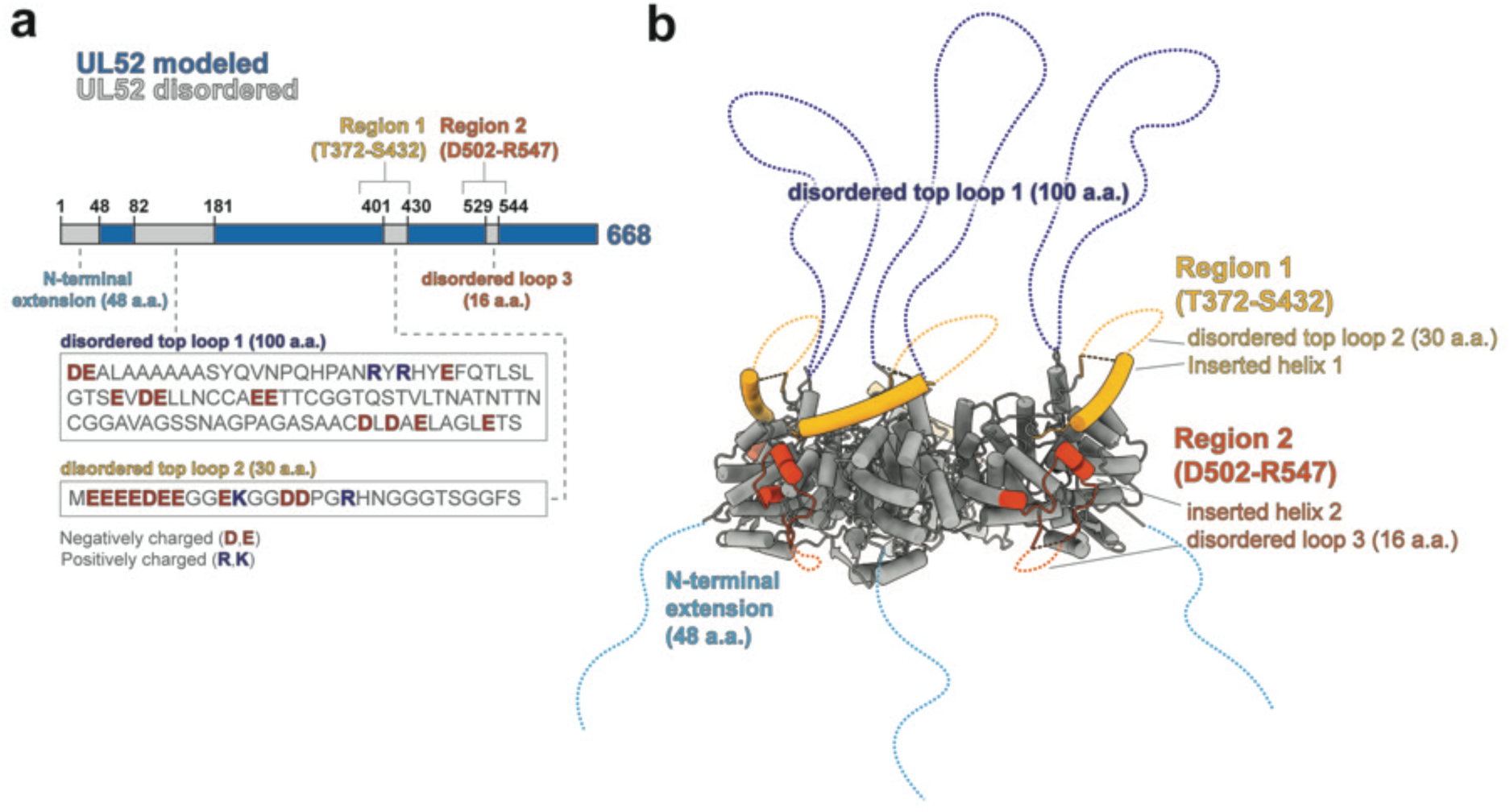
HCMV UL52 differs relative to ORF68. (**a**) Schematic of the UL52 sequence showing surface regions that could not be modeled (grey). Insets show the sequence of negatively charged disordered loops that extend from the top of the incomplete ring. (**b**) Model showing two major regions of UL52 surface that differ relative to ORF68 (yellow and orange). Unmodeled N-terminal extension (light blue) and long disordered top loop (dark blue) are represented as dashed lines.

**S4 Fig.**
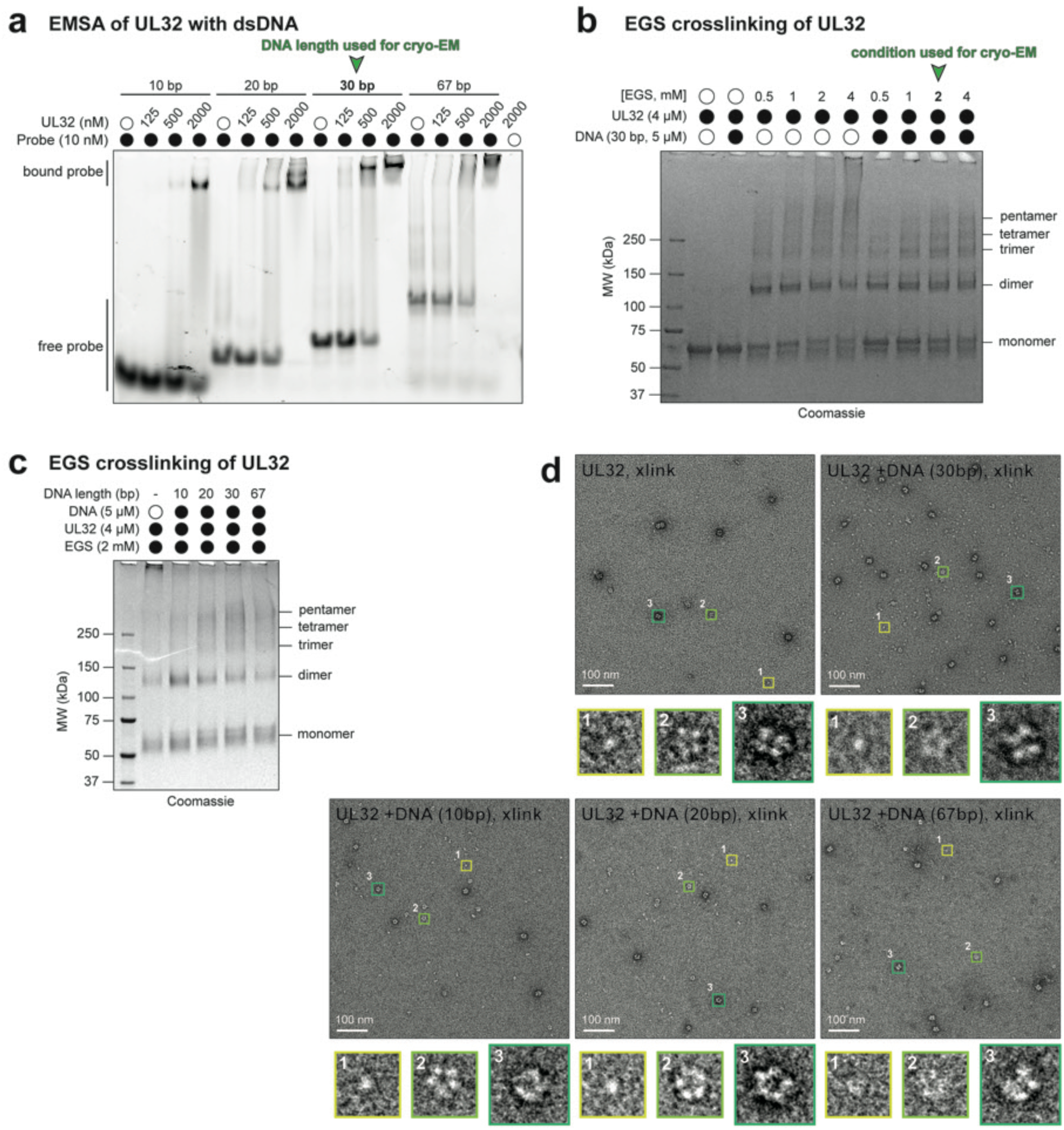
HSV-1 UL32 forms an unstable pentameric ring that can be stabilized by crosslinking. (**a**) Native gel for the electrophoretic mobility shift assay of UL32 binding to 5’-fluorescein labeled dsDNA probe of 10, 20, 30, or 67 bp. DNA probe used in cryo-EM sample (30 bp) is marked with an arrow. (**b**) SDS-PAGE gel showing UL32 crosslinked with and without DNA at EGS concentrations of 0.5 to 4 mM. The crosslinking condition used to prepare cryo-EM sample is marked with an arrow. (**c**) SDS-PAGE gel showing UL32 crosslinked with and without DNA of different lengths (10-67 bp). (**d**) Negative stain of UL32 crosslinked with EGS with and without DNA of different lengths (10-67 bp). Boxes indicate monomer (1, yellow), pentamer (2, light green), and higher-order species (3, green).

**S5 Fig.**
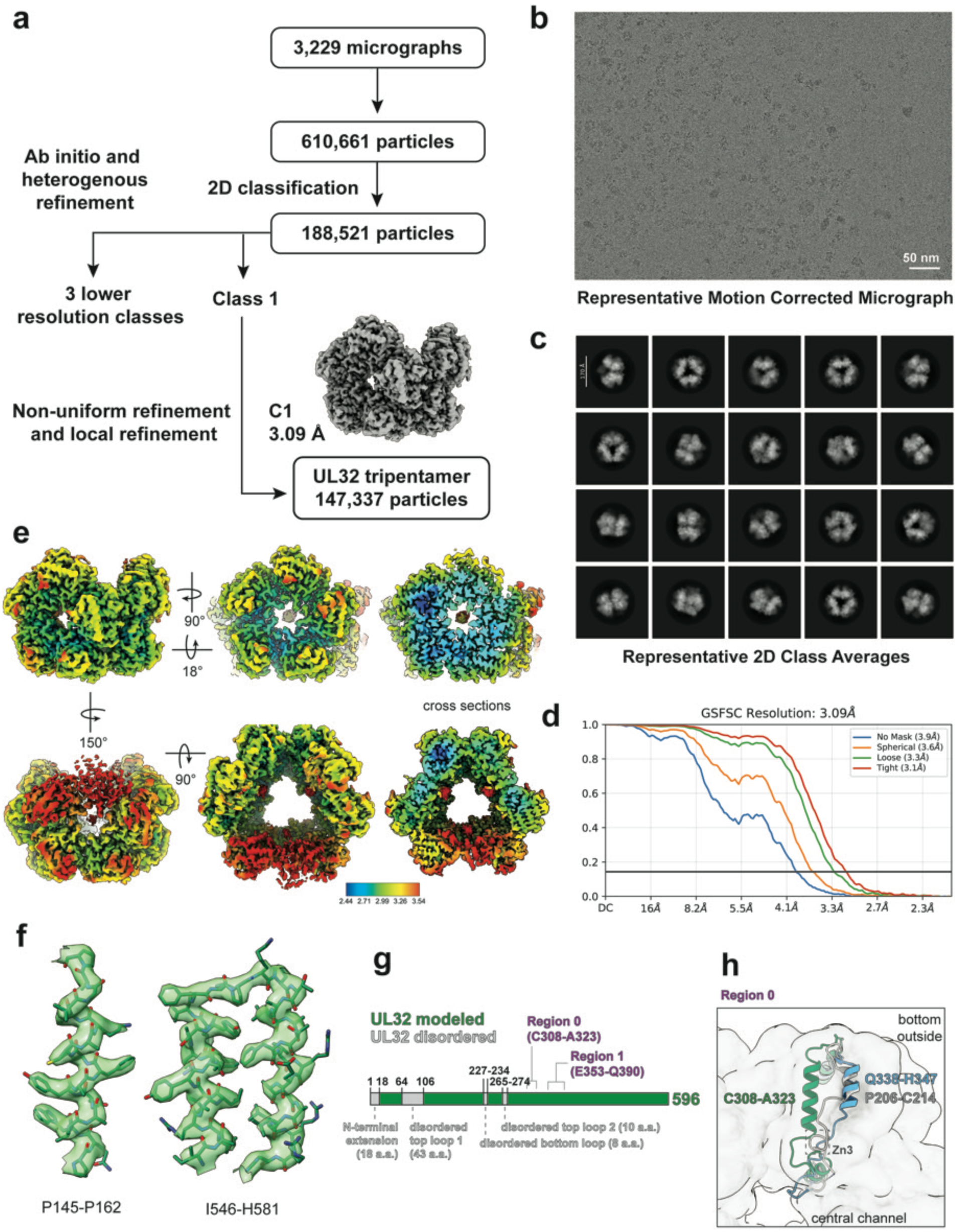
Cryo-EM data analysis for HSV-1 UL32. (**a**) Overall data processing workflow for the HSV-1 UL32 consensus cryo-EM reconstruction. (**b-c**) Representative micrograph with scale bar (50 nm) and 2D classes. (**d**) Gold-standard Fourier Shall Correlation (GS-FSC) curve for UL32 consensus map. (**e**) Local resolution for the UL32 consensus map. (**f**) Representative examples for the cryo-EM map and the corresponding modeled regions. Map has transparent surface; model is shown with sticks and heteroatom coloring. (**g**) Schematic of the UL32 sequence showing surface regions that could not be modeled (grey). (**h**) Bottom surface helix of UL32 (green) shown relative to UL52 and ORF68 (blue, grey). Model segments are shown inside transparent UL32 model surface at 10 Å resolution.

**S6 Fig.**
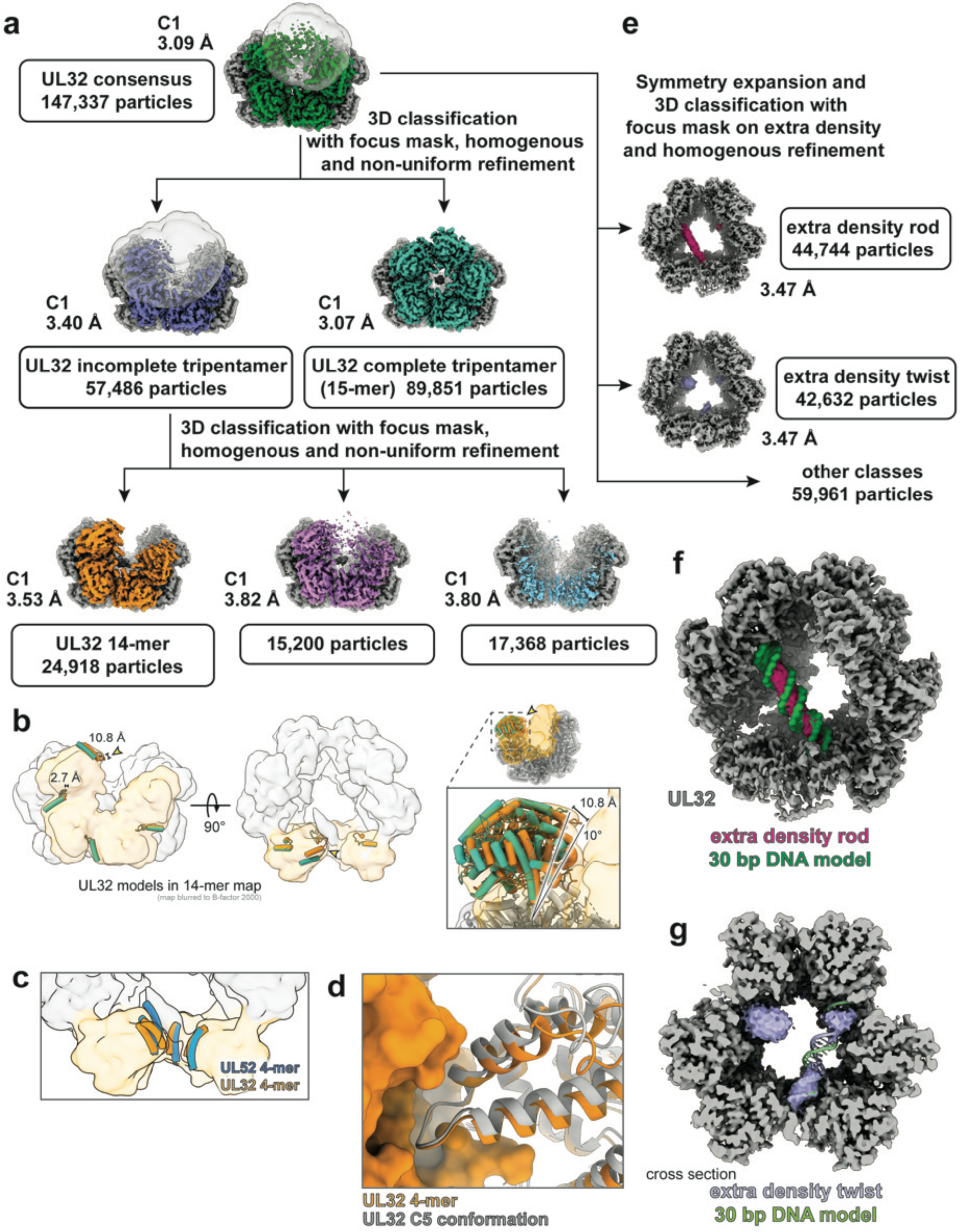
Further classification of UL32 consensus map. (**a**) Further 3D classification of the UL32 consensus cryo-EM density with focus masks on weak density regions of the third pentamer shows different occupancy of the third pentamer. Third pentamer density shown in various colors, first and second pentamer densities shown in grey, focus masks shown as transparent surfaces. (**b**) Pentamer interface loop and adjoining helix model segments (residues 448-468) for the UL32 4-membered incomplete ring (shown in orange) and pentamer (teal). Model segments shown inside the UL32 14-mer map (blurred to B-factor 2000 and shown as a transparent surface), with complete pentamers in grey and incomplete pentamers in orange. Inset: the most twisted protomer in the UL32 4-mer relative to UL32 pentamer is rotated by ∼10°. (**c**) Central channel helices of UL32 4-mer (orange) and UL52 4-mer models (blue). (**d**) Model segments shown inside the UL32 14-mer map (blurred to B-factor 2000 and shown as a transparent surface) with complete pentamers in grey and 4-membered incomplete ring in orange. (**e**) Symmetry expansion and further 3D classification of the UL32 consensus cryo-EM density with a focus mask on the extra density in the inter-pentamer space. UL32 density appears in grey, additional density appears in magenta and lilac. (**f-g**) UL32 classes from **e** modeled with 30 bp dsDNA model generated by AlphaFold3. **f**, DNA model shown as a molecular surface at 10 Å resolution (bright green) that fits the rod-like extra density shown in magenta. **g**, DNA model shown as a cartoon (green and lilac) that fits the twisted extra density shown in lilac. Density shown in cross section.

**S7 Fig.**
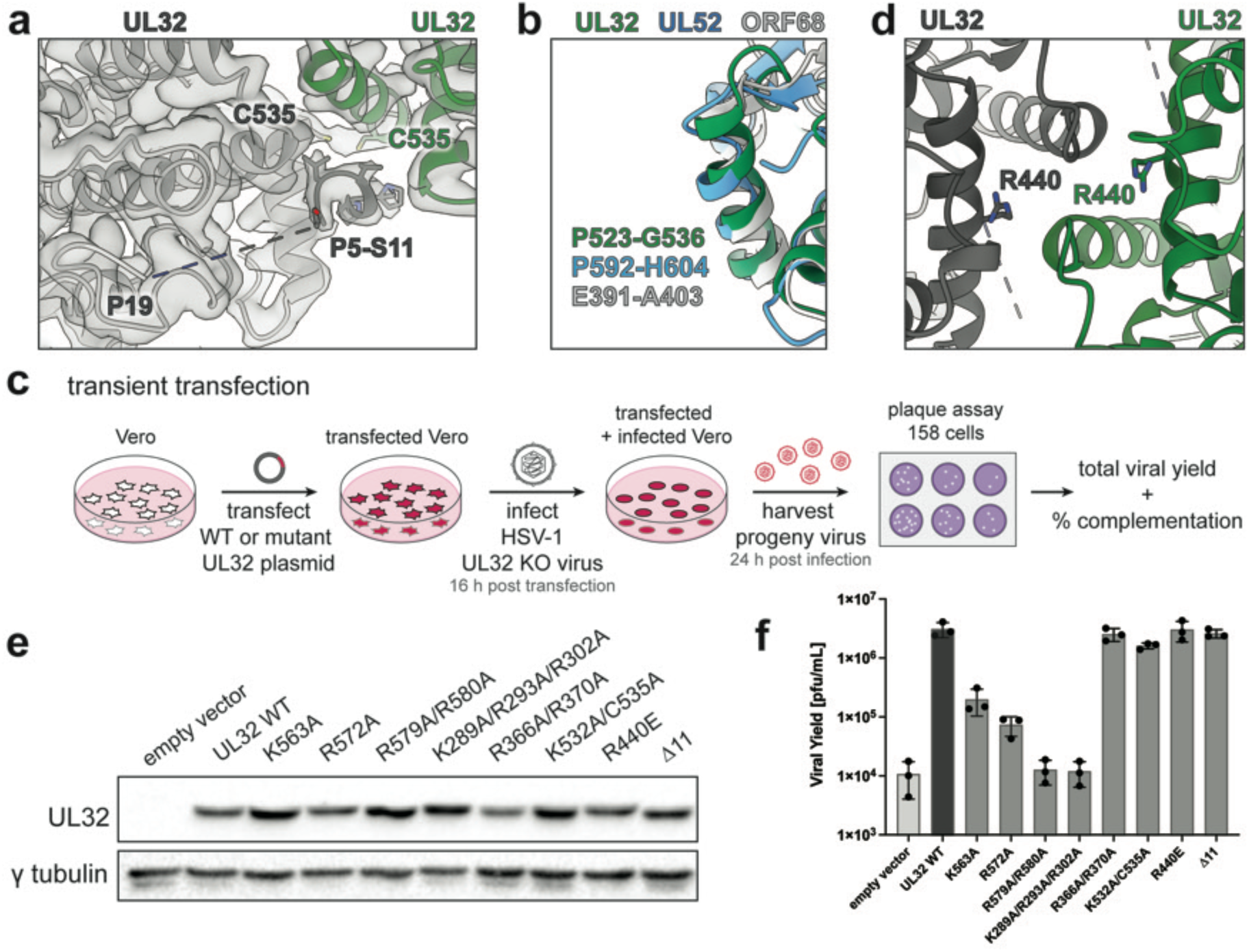
UL32 tripentamer interface and transient transfection assay. (**a**) Cryo-EM density map (transparent, moderate contour level) and model (cartoon, colored as in **Fig 4a**) of the UL32 tripentamer interface helix and additional stabilizing density modeled as P5-S11 of the N-terminal extension. (**b**) UL32 tripentamer interface helix (green) and corresponding helices from UL52 (blue) and ORF68 (grey). (**c**) Schematic of transient complementation assay. (**d**) Model of tripentamer interface highlighting charge swap mutant R440E. Model displayed colored as in **Fig 4a**. (**e**) Representative western blot for expression of UL32 WT and mutants in transient complementation assay. (**f**) Graph of total viral yield of UL32 mutants in a transient complementation assay from three independent biological replicates, used to calculate complementation efficiency in **Figs 4b** and **6c**. Error bars represent SD.

## SUPPLEMENTARY INFORMATION LEGENDS

**S1 Movie.** UL52 maps and model. UL52 3-mer map fades into 4-mer map, followed by fade into UL52 4-mer model. Detail of protomer interface is shown from the side. Colored as in Fig. 2.

**S2 Movie.** UL32 map and model. UL32 tripentamer map spins around 3-fold symmetry axis, then rotates to show top and bottom views along the 3-fold axis. Map fades into tripentamer model shown in tube helices, then rotates to view the pentamer model, shown in ribbons, along the local 5-fold symmetry axis. Detail of protomer interface is shown from the side. Colored as in Fig 3.

**S1 Raw Gel.** Original images for blots and gels in Figures 1B, 5C, Supplementary Figures S4A, S4B, S4C, and S7E.

## SUPPLEMENTARY INFORMATION 1

### Supporting Information - Original images for blots and gels

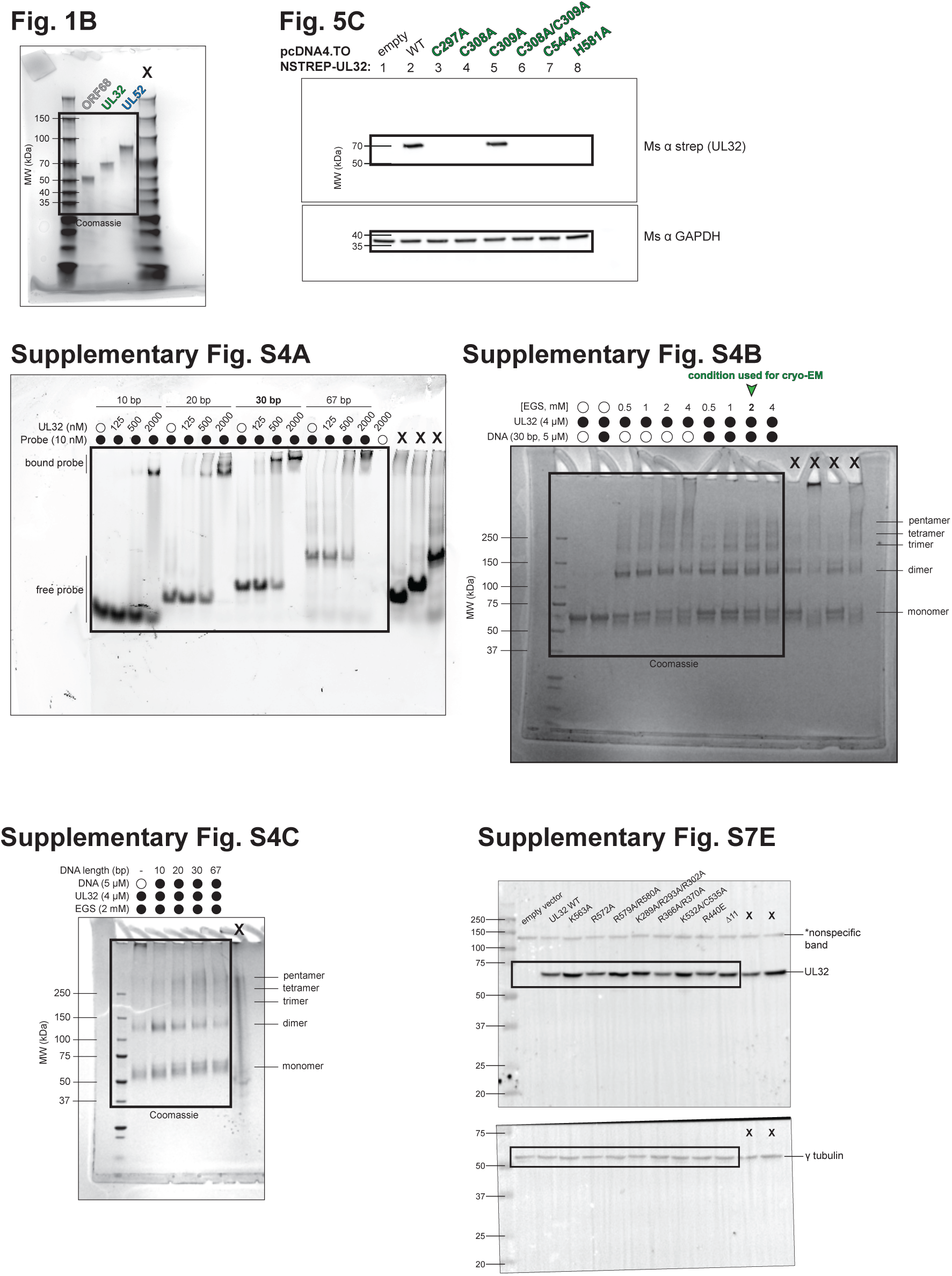

## Notes

### Competing Interest Statement

The authors have declared no competing interest.

### Summary of Updates

Updated based on reviewer feedback from Review Commons and Plos Pathogens. Reanalyzed 3mer/4mer dataset to better separate particles, clarification about expression system and result on oligomeric state, additional discussion of alternative possibilities for biological role of oligomeric states, additional statistical analyses, modification of language for structural differences throughout the manuscript.

